# Transcription factor handover reshapes hub dynamics to coordinate temporal progression of Notch response

**DOI:** 10.64898/2026.11.11.765832

**Authors:** Carmen Santa Cruz Mateos, Charalambos Roussos, Angelette Mendonca, Sarah J Bray

**Author notes:** co-corresponding authors: CSCM,; SJB.

## Abstract

To decipher information in space and time, cells require mechanisms to sustain gene expression programmes once an initial signal decays. Localised condensation of transcription complexes into hubs is emerging as a key step in gene regulation and, by combining live imaging of transcription and of transcription complexes at target loci, we have investigated how these evolve to enable a prolonged transcription outcome in response to Notch signalling. Measuring the dynamics of *hindsight* (*hnt*) transcription reveals two phases. The first, a probabilistic response similar to *E(spl)-C* genes, is directly controlled by Notch and the second, a sustained response, is indirectly controlled through the BTB transcription factor Broad. The two phases correlate with a transition in transcription factor enrichment at *hnt*: complexes of Notch coactivator Mastermind and of Broad are successively enriched. Notably, the lifetime of the hubs formed by each are distinct and correlate with the change in transcription profiles. These reflect the binding dynamics of each complex, with Broad having significantly more stable association with chromatin, forming longer lasting hubs and yielding sustained transcription. Furthermore at intermediate stages we detect hubs containing both Broad and Mam implying an evolution in hub composition. Together our data support that the dynamic properties of transcription hubs are crucial for the probability and dynamics of transcription and point towards a “hub handover” in enabling temporal progression in gene expression responses.

## Introduction

To properly differentiate and function, cells must integrate and decipher multiple signals in space and time to bring about appropriate gene expression responses. Failure to do so correctly can lead to severe developmental defects and diseases^1–3^. While the mechanisms that control the immediate response to an incoming signal are becoming better understood along with the role of combinatorial factors in conferring precise spatial gene activity, it remains unclear how transcription of genes is regulated in time once the initial signal decays. In part this is because gene expression dynamics have primarily been studied during short developmental proceses^4–7^. Therefore, there is a need to understand how cells mediate prolonged transcription responses in time during differentiation processes.

Notch signalling is one of the highly conserved pathways that is fundamental for cell fate acquisition, stem cell maintenance and tissue function where it generates diverse transcriptional outcomes^8–12^. Binding of the cell-surface Delta ligand to the Notch receptor provokes receptor activation and cleavage, liberating the intracellular domain, NICD. In the nucleus, the released NICD forms a complex with the DNA-binding transcription factor CSL (Su(H) in flies) and the co-activator Mastermind (Mam) to promote transcription of target genes, notably those of the HES gene family^13–15^. In the tissues where this has been examined, successful target gene activation involves the concentration of NICD-Mam-CSL transcription complexes into localised transcription factor hubs which precede and inform the levels of transcription^16–18^. Whether this extends more widely and to other loci besides the well-characterised multi-gene *Enhancer of split-Complex* in *Drosophila* [*E(spl)-C*] remains to be established. Furthermore, it is unclear what mechanisms operate to sustain target gene transcription once Notch activity declines.

The ability to visualise and track transcription factors (TFs) in vivo has revealed that they form localized condensates or hubs that concentrate key components of the transcription machinery and enable enhancer-promoter cross-talk to drive bursts of transcription^16–24^. Bursts are generated by periods of promoter activity interspaced by periods of inactivity^25,26^ and are regulated by enhancers^7,27,28^, suggesting that enhancer-promoter communication is dynamic rather than static. In many developmental contexts, bursts are comparatively short-lived, raising the question how longer lasting responses are generated. Furthermore, during lineage acquisition, the regulation of gene activity in time and space often involves a switch in enhancer usage, so-called “handover” model^29,30^, which relies on temporal segregation of TF expression^31,32^. It remains to be established how these complex temporal requirements can be reconciled with a model involving transient transcription hubs as key intermediaries.

To investigate how transcription factor hubs regulate transcription outcome in time, we focussed on the *Drosophila* ovary, where Notch signalling is endogenously active for an extended period^33–35^. Two direct target loci have been identified, *E(spl)-C* and *hindsight* (*hnt*) genes^33–35^. Notably, their expression profiles differ with *hnt* expression being considerably more prolonged, with an activity period that extends for over 25 hours and is required for cells to enter in endocycle^35–37^. By characterising the transcriptional dynamics of the different target loci, using the MS2/MCP system and smFISH, together with simultaneous tracking the behaviour of TFs by real time imaging in relation to the cognate target locus, we found that *hnt* undergoes two dynamically different phases of transcription. Each phase is controlled by a different transcription factor complex, namely those of Notch and Broad, which are successively enriched in a hub at the *hnt* locus . Furthermore, the lifetime of the hub and the properties of the two transcription complexes relate to the transcription profiles at each stage. Overall, our data invoke a direct role for TF hubs in coordinating the dynamics of transcription over time and demonstrate how a handover between different TF complexes enables the evolution of a prolonged transcription response to support a developmental outcome.

## Results

### Expression of *hindsight (hnt)* extends beyond Notch activity

In the *Drosophila* follicular epithelium (FE) Notch activity controls transcription of several different target genes whose precise temporal profiles appear to differ across a prolonged developmental window^33,35–37^. To elucidate the fundamental principles governing the differing responses, we focused our analysis on two primary downstream target loci: *E(spl)-C* genes and *hindsight* (*hnt)* (Figure 1A). We utilized single-molecule fluorescence in situ hybridization (smFISH) to monitor active transcription sites (ATSs), manifest as intense, high-density puncta within the nucleoplasm (Figure 1B). The proportion of follicle cell (FC) nuclei containing an active ATS provided a quantitative measure of transcriptional activity at each developmental stage^38^. At earlier stages (S5-S7), the proportion actively transcribing nuclei was comparable between the different target genes (Figure 1C). However, they diverged at later stages (S8, S9) (Figure 1B, C). The proportion of nuclei actively transcribing *E(spl)-C* genes decreased substantially by S9 whereas the converse was true for *hnt*, with circa 80% of nuclei containing an *hnt* ATS at S9 (Figure 1B, C). Comparing the two directly through stages 5-9 emphasises that *hnt* expression has an increasingly upward trajectory in relation to that of *E(spl)-C* (Figure S1A).

**Fig. 1.**
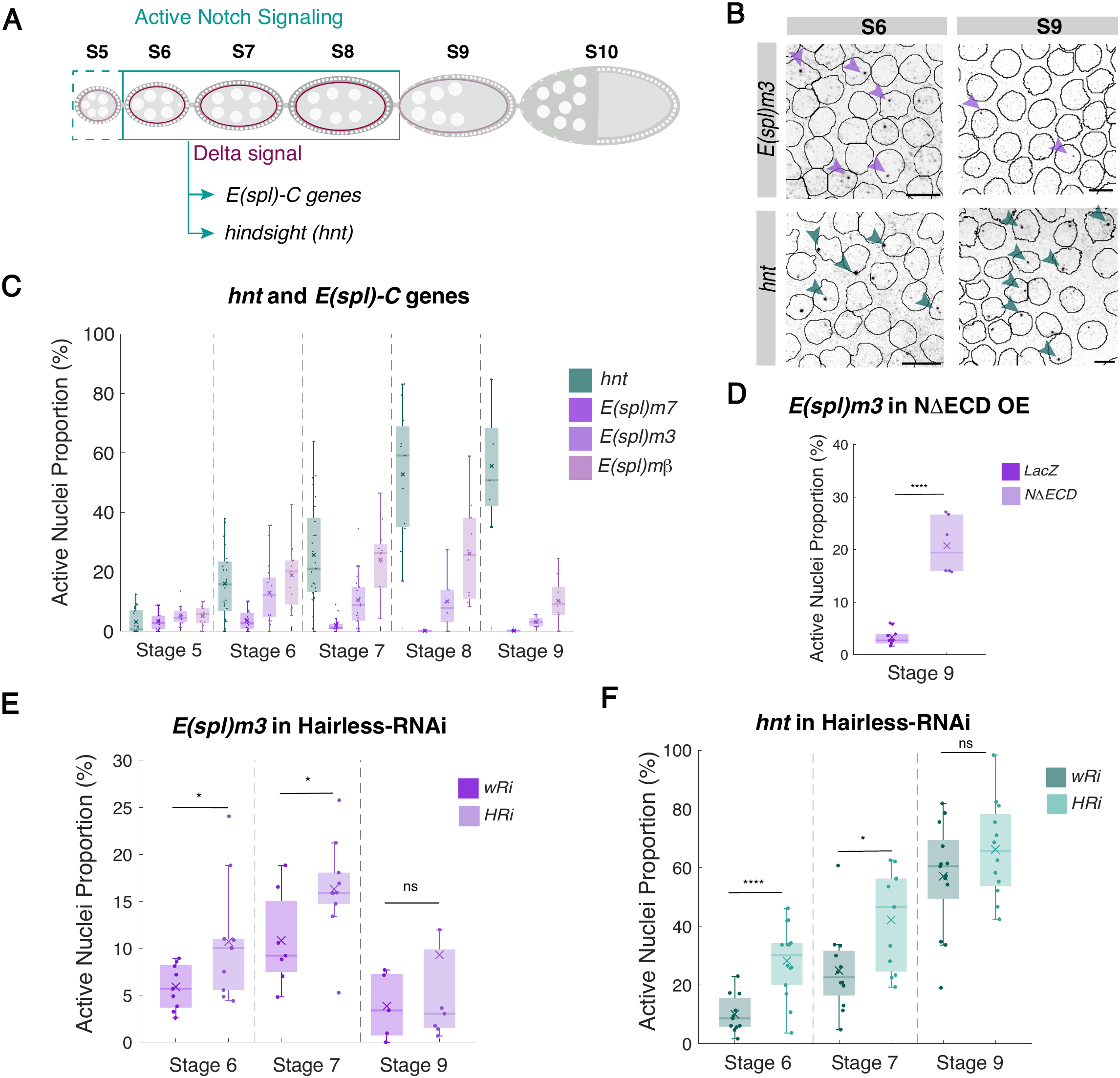
*hnt* expression diverges from Notch activity at late stages. **(A)** Cartoon of stage(S) 5-10 egg chambers (e.c.) from a *Drosophila* ovariole, FE monolayer surrounds germline cells (large nuclei) which express Delta (magenta) leading to Notch activity in FE during S6-9 (turquoise square). Dashed line indicates start of Notch signal at stage 5. **(B)** Confocal images of smFISH with *E(spl)m3*-570 and *hnt*-670 probes in wild-type stage 6 and 9, with segmented nuclei outlined. ATS are indicated (arrowheads). Scale bars, 5μm. **(C)** Proportions of nuclei actively transcribing E*(spl)-C* genes and *hnt* at indicated stages. Mean data per e.c.; n = 4 to 27.**(D)** Proportions of nuclei with *E(spl)m3* ATS in control (*LacZ*) and NΔECD expression conditions detected by smFISH at stage 9. Mean data per e.c.; n = 6. **(E)** Proportions of FE nuclei with *E(spl)m3* ATS in control (*white-RNAi, wRi*) and *Hairless RNAi* (*HRi*) at stages indicated. Mean data per e.c.; n = 7 to 10 **(F)** Proportions of FE nuclei with hnt ATS in control (*wRi*) and Hairless RNAi *(HRi*) at stages indicated. Mean data per e.c.; n =10 to 14.

The profile of *E(spl)-C* gene activity closely mirrored the predicted pattern of Notch activity, based on the decay of germline Delta expression by S9 (Figure 1A, C). If Notch activity levels are the defining feature for *E(spl)-C* transcription, their expression should be prolonged in conditions that extend Notch activity. To test this prediction, we expressed a constitutively active form of Notch, NotchΔECD, throughout the FE and monitored the consequences on E*(spl)-m3* transcription at S9 when it has normally declined. In agreement with the model that decline in Notch activity normally limits its transcription, the proportion of nuclei with E*(spl)-m3* ATS at S9 was substantially elevated in ovarioles with NotchΔECD (Figure 1D, S1B). As a complementary strategy, we tested whether downregulating the core co-repressor Hairless had an impact on *E(spl)-m3* transcription. Reduced co-repressor levels are expected to elevate the amounts of active Notch transcription complexes in nuclei where the pathway is active. Depletion of Hairless, using a previously validated *Hairless*-RNAi treatment^17,39^, increased the proportion of nuclei with *E(spl)-m3 ATS* at earlier stages (S6,S7) but not in S9, consistent with inferred decline in Notch activity during S9 (Figure 1E, S1C,D).

Unlike *E(spl)m3*, *hnt* transcriptional activity increased after S7, achieving highest levels at S9. This suggests that *hnt* expression might decouple from Notch pathway at this later stage. If this is the case, *hnt* expression levels should only be sensitive to Hairless depletion at earlier stages and not at S9. In agreement, the proportion of nuclei actively transcribing *hnt* was markedly enhanced by Hairless depletion during S6 and S7, (Figure 1,F; Figure S1E,F) but was not significantly altered at S9. On the basis of these findings, we propose that two distinct regulatory regimes coordinate the transcriptional response of *hnt*: an early phase (S5–7) where *hnt* transcription is dependent on Notch signalling and a second phase where *hnt* response becomes Notch independent and is orchestrated by another factor.

### *hnt* transcription displays two distinct dynamic phases

The smFISH analysis indicates two phases of *hnt* transcription with only the first being directly dependent on Notch activity. To investigate further the properties of these two phases, we used the MS2/MCP system to track nascent transcription of *hnt* in real time^40^, focusing on S6 and S9. We used a strain carrying 24xMS2 loops within the *hnt* 5’ UTR^41^ (Figure 2A) and combined this with MCP::GFP to detect nascent transcription, and Histone::RFP to segment and track nuclei. Periods of active transcription are evident by the presence fluorescent nuclear puncta whose intensity levels are a direct read-out of RNA molecules produced (Figure 2B). Tracking these *hnt-MS2/MCP::GFP* (Figure 2C) puncta revealed that *hnt* transcription occurred sporadically with periods of activity (Active periods) interspersed with periods of inactivity (Inactive periods) that were asynchronous among follicle cells (Figure 2C). At S6, a relatively small proportion-20% - of follicle cells were transcribing *hnt* at any given time point (Figure 2C,D). By contrast, at S9 a substantially higher proportion of cells were active at any time, averaging close to 50% with up to 70% transcribing in some egg chambers (Figure 2C,D). These proportions are comparable to those detected by smFISH at equivalent time points (Figure 1C).

**Fig. 2.**
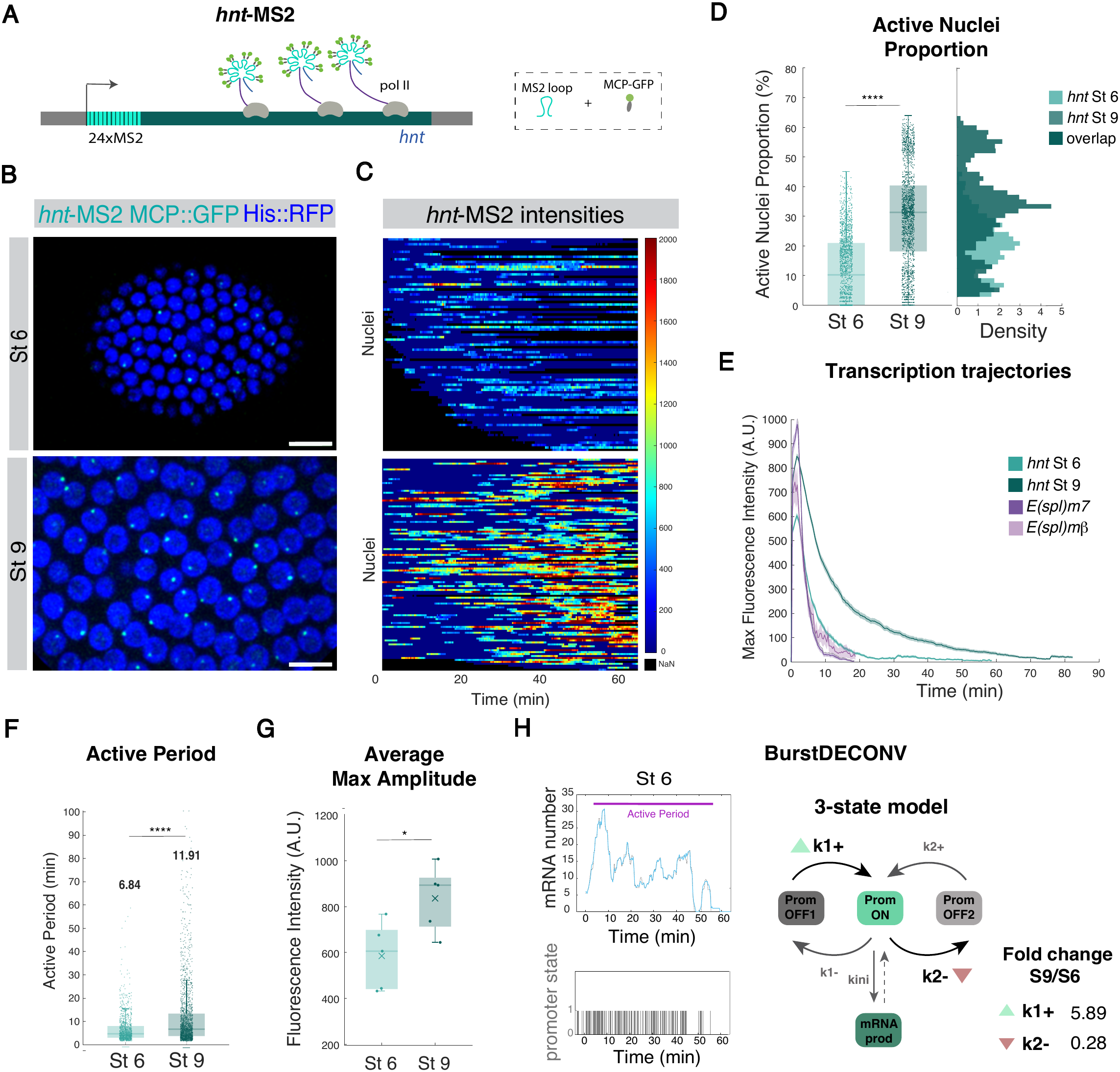
Dynamics of *hnt* transcription shows two different phases. **(A)** Diagram illustrating the MS2/MCP system. 24x MS2 loops (cyan) have been introduced into 5’UTR of *hnt* (teal) coding regions^29^. MCP::GFP (green) binds to MS2 loops in nascent mRNA (Pol II; grey). **(B)** Images from in vivo movies of *hnt-*MS2/MCP::GFP, transcription puncta (cyan) are detected within nuclei (blue; h2Av-RFP), stages as indicated. Scale bars, 10 μm. **(C)** Heatmaps of maximum fluorescence intensity of *hnt*-MS2/MCP::GFP in active nuclei over time at stage 6 and stage 9 (Movies S1 and S2). **(D)** Average proportion of nuclei transcribing *hnt*-MS2 per time frame (20 seconds) at stage 6 (turquoise) vs stage 9 (teal). Probability density function (pdf) of frames (20 seconds) containing indicated proportions of nuclei transcribing *hnt*-MS2/MCP::GFP at stage 6 (turquoise) vs stage 9 (teal). Darker teal represents overlapping numbers between the two stages. p=1.1353E-118. **(E)**Average intensities of *hnt*-MS2 traces at stage 6 (turquoise) vs stage 9 (teal) together with traces of *E(spl)m7*-MS2 (purple) and *E(spl)mβ*-MS2 (lilac) at stage 6 for comparison. Shading represents the SEM. *hnt*-MS2 St 6 (n = 993 nuclei) and St 9 (n = 1644 nuclei) from at least 5 e.c. **(F)** Active period durations for *hnt*-MS2 (at stage 6 (turquoise) vs stage 9 (teal). Mean values are indicated above data. p=1.30163E-25. **(G)** Average Intensities during the first 3 minutes of *hnt*-MS2 transcripion at stage 6 (turquoise) vs stage 9 (teal). (D, E and G) Boxplots indicate the median, with 25 to 75 quartiles; error bars are SD. **(H)** Example of MS2/MCP::GFP trace from *hnt* transcription at St 6, with calibrated mRNA numbers through time (upper pannel) and inferred Pol II initiation events calculated using BurstDECONV (bottom pannel). A 3 state model that illustrates the steps controlled by each rate. Green triangle indicates an increase in k1+ rate fold-change and red triangle a decrease in k2-, Fold change values of the obtained rates between St6 and St9 are presented. k1+ and k2-differ between stage 6 and 9, resulting in longer burst durations in the latter.

Other features of *hnt* transcription also differed between S6 and S9. At S6 *hnt* transcription profiles were similar to those observed for *E(spl)-C* genes at the same stage (*E(spl)-m7* and *E(spl)-mβ)* ^18^, with low proportions of active nuclei and comparable Active Periods (Figure 2D-F, Figure S2A). This supports the hypothesis that the *hnt* response is directed by Notch signalling at this stage. At S9, *hnt* profiles consisted of substantially longer Active Periods whose average duration was almost double (11.91 min) that observed at S6 (6.84 min) (Figure 2F). As evident when transcription traces were aligned (Figure 2E), many nuclei (16%) had Active Periods longer than 20 minutes, with some exceeding one hour (Figure 2E, S2A). Comparison of the amplitude of the MS2/MCP traces at both stages, a direct readout of the number of nascent mRNA molecules, revealed that the transcriptional output was also increased at S9 (Figure 2E,G).

Gene transcription profiles are the result of multiple ON and OFF cycles of promoter activity that confer transcriptional bursts^25,26^. To analyse the promoter dynamics of the *hnt* gene during the different phases, we used a statistical inference method, BurstDECONV, to resolve the MS2/MCP signal traces into transcription initiation events^25,26^ (Figure 2H, S2B). This generates a distribution of waiting times between successive polymerase initiation events. The number of promoter states is then inferred by the minimum number of exponentials required to fit the survival curve of these waiting times. In the case of *hnt* this was three, suggesting its expression profile drives from three promoter states (Figure 2H): a competent promoter-ON state and two distinct promoter-OFF states of different lengths. The latter may be explained by the existence of a longer polymerase paused state, as seen in another developmental contexts^42^. The main differences between *hnt* transcription kinetics at stage 6 and 9 were a 5-fold increase in k1+, the rate at which the promoter leaves the longer OFF to enter the ON state, and a five fold decrease in k2-, the rate from ON state to the shorter OFF (Figure 2H, S2C). These two different rates account for *hnt* having longer burst ON durations and higher burst frequency at the later stage ^25,26^. *hnt* transcription thus undergoes two phases that differ in their dynamics, the first being more transient and probabilistic and the second more sustained and penetrant brought about by changes in burst duractions and frequency.

### The two phases of the *hnt* response are orchestrated principally by Notch and the zinc-finger transcription factor Broad

The most likely explanation for the two distinct phases in *hnt* transcription dynamics, is that they are coordinated by different enhancers, only one of which is directly regulated by Notch. Three Notch-responsive enhancers were previously identified within *hnt,* based on regions bound by CSL in blood cells^43^ (Figure 3A). Reporters comprising these enhancers upstream of a minimal promoter and coding sequences for eGFP are expressed in a Notch dependant manner in various tissues (Figure 3A) ^43^. We therefore examined their expression in the ovary and, of the three, only *hnt^peb-3^* was active in the FE from S5 to S8 (Figure 3B, S3A). The *hnt^peb-3^* expression window coincides precisely with the proposed Notch regulated period of *hnt*, favouring the model that *hnt* is a direct target of Notch activity during this phase.

**Fig. 3.**
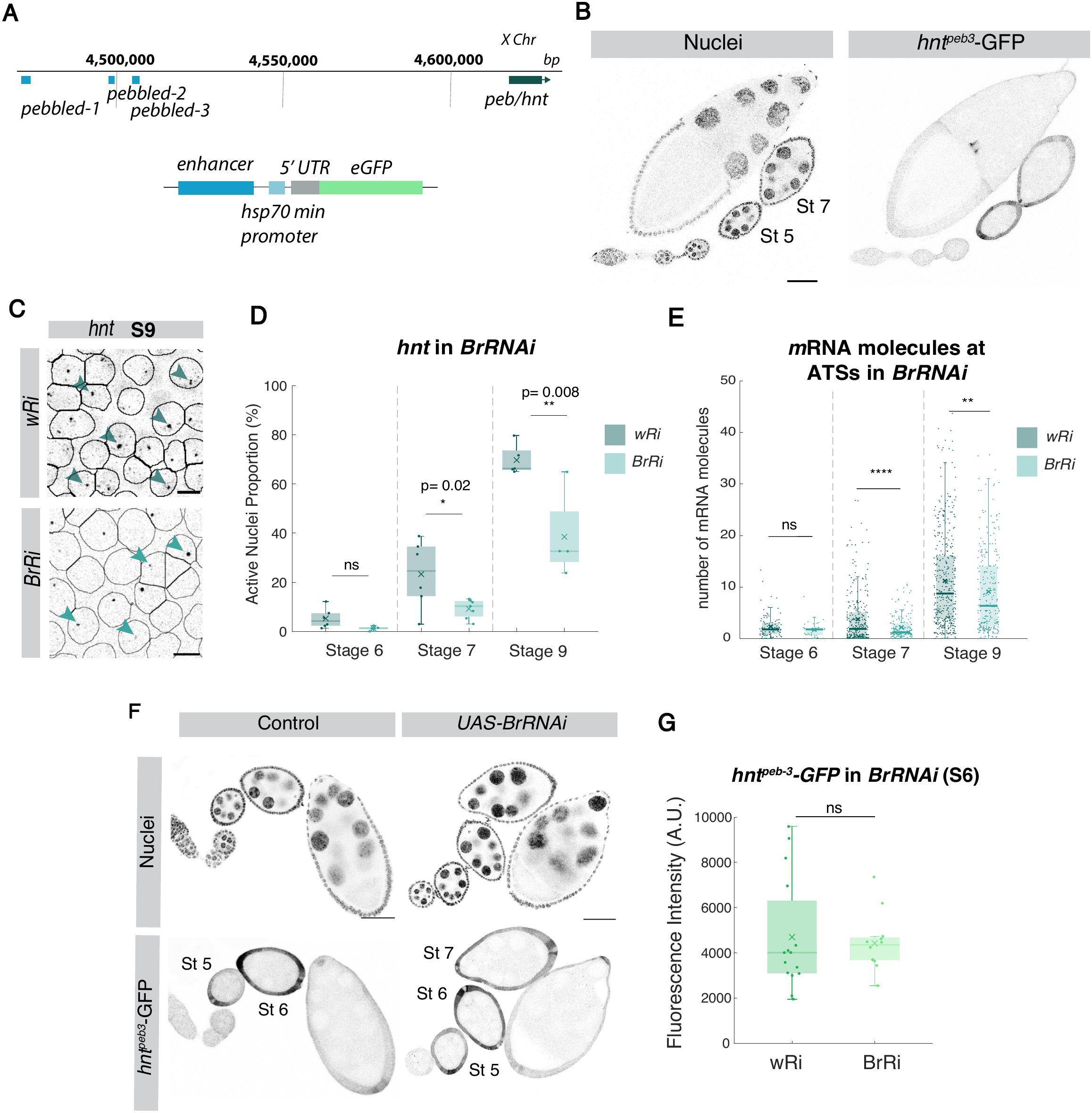
Notch and TF Broad direct first and second phases of *hnt* transcription respectively. **(A)** Diagram showing location of the three Notch responsive enhancers found in *hnt* upstream regulatory region, coordinates X Chromosome dm6, with schematic of enhancer reporters^43^. **(B)** Confocal images from an ovariole; nuclei detected with DAPI and corresponding *hnt^peb-3^*-GFP expression, present from stage 5 to 7. **(C)** Confocal images of smFISH, *hnt*-670 probes in control (*wRi*) and Broad-RNAi (*BrRi*) at S9 with masks from nuclei segmentation superposed. Scale bar, 5 μm. **(D)** Proportion of nuclei with *hnt* ATS detected by smFISH in control (*wRi*) and Br-RNAi (*BrRi*) conditions at stages indicated. (n=4-7 e.c). **(E)** Number of molecules at *hnt* ATSs in control (*wRi*) and Br-RNAi (*BrRi*) conditions at stages indicated. n=102, 35, 294, 179, 418, 279 from 4-7 e.c. **(F)** Expression of *hnt^peb-3^-GFP* in control (*wRi)* and Broad-RNAi (*BrRi*) conditions. **(G)** Quantification of *hnt^peb-3^-GFP* levels at S6 in *wRi* and *BrRi* conditions. *hnt* locus Halo::Mam *hnt* locus BrZ1::GFP

The second phase of *hnt* transcription begins during S7 and persists through S9. To identify the factors regulating *hnt* activation during this period, we first selected candidate transcription factors whose expression coincided with this time window based on single cell RNA sequencing data^44,45^. Primary candidates included the homeodomain transcription factor Mirror and the zinc-finger transcription factor Broad^46^. We therefore tested whether they contribute to *hnt* regulation by using validated RNAi lines to deplete their expression levels in FE (see Methods) (Figure S3B) and measuring the effects on the proportion of nuclei with *hnt* ATS. Depletion of Broad^47,48^ (RNAi targeting all isoforms) substantially reduced the proportion of nuclei with *hnt* ATS at S7 and S9 (Figure 3C,D) whereas depletion of Mirror had no significant effect at either stage (Figure S3D). Furthermore, Broad depletion also led to a decrease in ATS average fluorescence intensity at S7 and S9, indicating that, in those nuclei that retained activity, fewer RNA molecules were produced. (Figure 3E). In contrast, Broad knockdown had no affect on transcription of *E(spl)m7* (Figure S3C), nor did it affect expression of *hnt^peb-3^* reporter (Figure 3F,G). Thus, these data support a model where Broad takes over the regulation of *hnt* from S7, acting through a distinct enhancer(s).

### Mastermind and Broad enrichment at *hnt* corresponds temporally to their regulatory phase

In response to Notch activity, the co-activator Mastermind (Mam), together with its partners in the Notch activation complex, becomes enriched at *E(spl)-C,* forming a transcription condensate or hub^17,18^. In FE these are present during S5-S7 in nuclei, detected by recruitment of endogenous Mam::Halo where the genes are transcriptionally active^18^. We therefore hypothesised that comparable Mam hubs would form at the *hnt* locus during the Notch-active period. To investigate we generated a strain in which the *hnt* locus could be detected by live imaging, using the ParB/Int system^49^. Using CRISPR engineering, the *Int* sequence was inserted into *hnt* in a non-conserved regulatory region that showed no CSL binding in embryonic ChIP-seq data (Figure 4A; see Materials and Methods), rendering the *hnt* locus visible in the nucleus when in combination with fluorescent ParB. As previously reported, Mam distribution is heterogeneous within FE nuclei (Figure 4B) and when examined in conjunction with the *hnt* locus tag, it was evident that dense Mam::Halo puncta co-localised with *hnt* (Figure 4C). Thus, as for *E(spl)-C*, Mam forms a concentrated hub at *hnt* in FE nuclei. This is notable because, *hnt* is a single gene locus, unlike *E(spl)-C* which contains several Notch responsive genes and hence might have unusual properties.

**Fig. 4.**
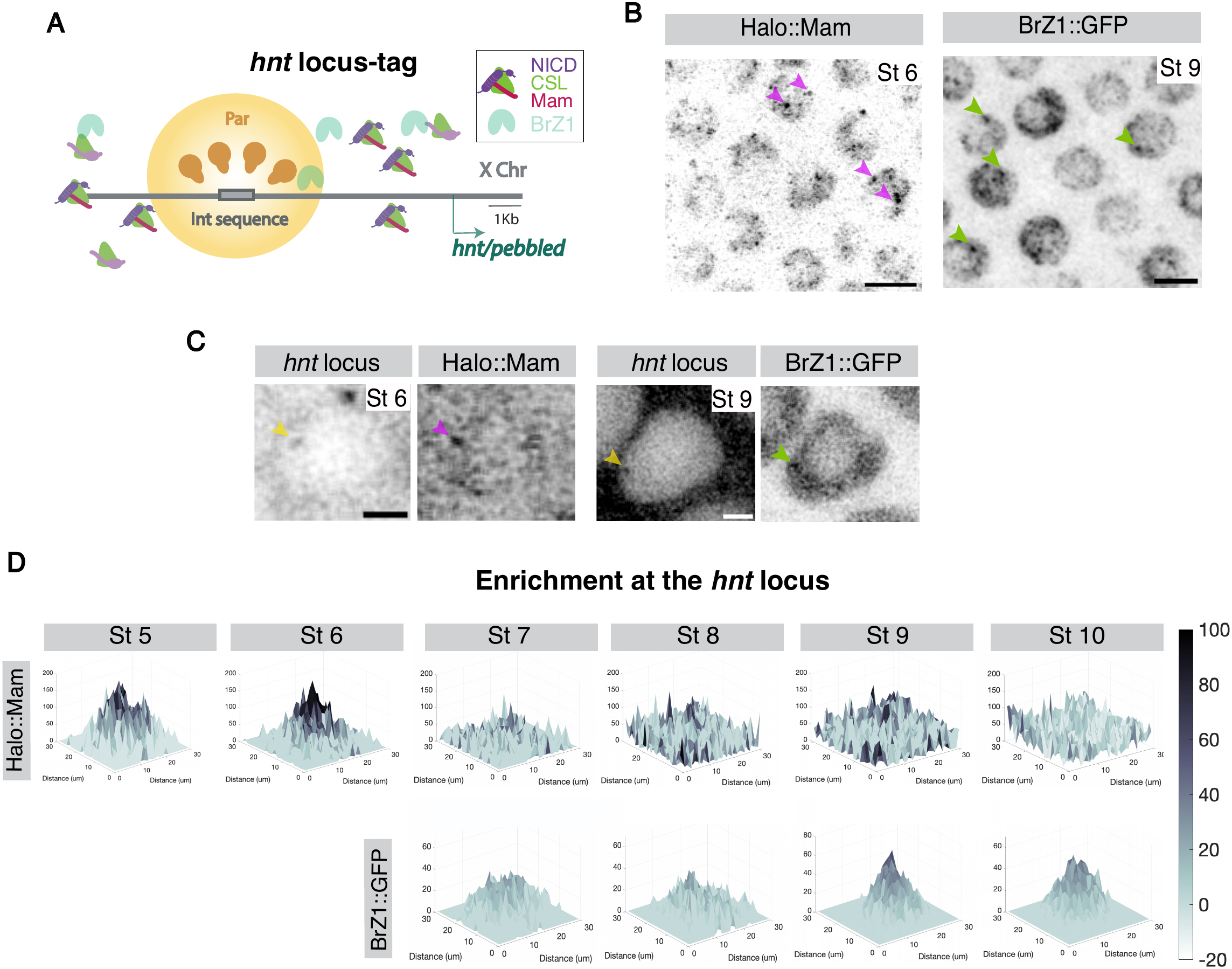
Both Mam and Broad form hubs at *hnt* locus. **(A)** Diagram summarizing Par/Int labelling of *hnt* locus. Par protein (yellow) is recruited to int sequence (gray) inserted within *hnt* locus. Co-recruitment of complexes [NICD (purple), CSL (green), and Mam (magenta)] or Broad-Z1 (light green) can be measured. **(B)** Confocal live images, Halo::Mam at S6 and BrZ1::GFP at S9. Magenta and green arrowheads indicate examples of putati-ve hubs. Scale bar, 5 μm. **(C)** Live imaging of individual FC nuclei stage 6 and stage with *hnt* locus (yellow arrowheads, stage 6 and stage 9 as indicated) and corresponding localized enrichment of Halo::Mam (magenta arrowhead) and BrZ1::GFP (green arrowhead). Scale bars represent 2 μm. **(D)** Normalized Mean Halo::Mam and BrZ1::GFP intensities centered at *hnt* locus throughout the stages indicated, visualized by 3D surface plots. n=110, 130, 85, 113, 111,100 from at least 3 e.c.for Mam and n=98, 92, 183, 214 from at least 3 e.c.for Broad.

We next asked whether the major Broad isoform, BrZ1, exhibited similar hub-forming behaviour. To address this, we used BrZ1::GFP and combined it with the ParB/Int tagged *hnt*. Live imaging of FE nuclei at S9 confirmed that Br-Z1 was heterogeneously distributed within the nucleoplasm, as for Mam, consistent with it forming condensates (Figure 4B). Importantly, puncta of BrZ1 enrichment colocalised with the *hnt* locus during S9 as predicted if it forms a transcription hub at this stage (Figure 4C).

Ǫuantifying enrichment of each factor at the tagged *hnt* at different stages revealed that the periods of Mam and BrZ1 recruitment were complementary (Figure 3B,D, 4D). Significant enrichment of Mam::Halo occurred predominantly at S5 and S6 (Figure 4D), with low levels persisting at S7 and S8 (Figure S4A). In contrast, the most prominent enrichment of BrZ1::GFP was S9 and S10, with lower levels present from S7 (Figure 4D). The complementary periods of Mam and BrZ1 recruitment thus coincide with the distinct phases of *hnt* regulation (Figure 3B,D, 4D) suggesting these hubs have a key role in coordinating the sequential phases of *hnt* transcription.

### Mam and Broad dynamics relate to those of concomitant transcription

The periods during which Mam and BrZ1 are enriched at *hnt* correspond to those with dynamically distinct phases of *hnt* transcription. As we have previously detected a relationship between Mam hub durations and *E(spl)-C* gene transcription dynamics, we hypothesised that the characteristics of the Mam and Br-Z1 hubs might correlate with the ensuing *hnt* transcription profiles. We first performed high resolution live imaging of Halo::Mam in combination with the ParB/Int-labelled *hnt* locus and *hnt*-MS2/MCP to visualize transcription (Figure 5A, Movie S3). Aligning the tracked Mam signal at *hnt* by the onset of transcription revealed that Mam levels peaked shortly (2.5 minutes) before transcription began (Figure 5B) as was observed at *E(spl)-C*^18^. Notably, the Mam hubs persisted for less than 5 minutes, a duration that coincided with that of the subsequent active period of *hnt* transcription (Figure 5B). When MS2 traces were displaced in time, cross-correlation analysis peaked at - 4 minutes, suggesting that Mam and MS2 traces correlate with this time delay (Figure S5A). These properties are similar to those observed at *E(spl)-C*, supporting the conclusion that Mam hubs are highly dynamic, and inform the subsequent transcription profile of the cognate target.

**Fig. 5.**
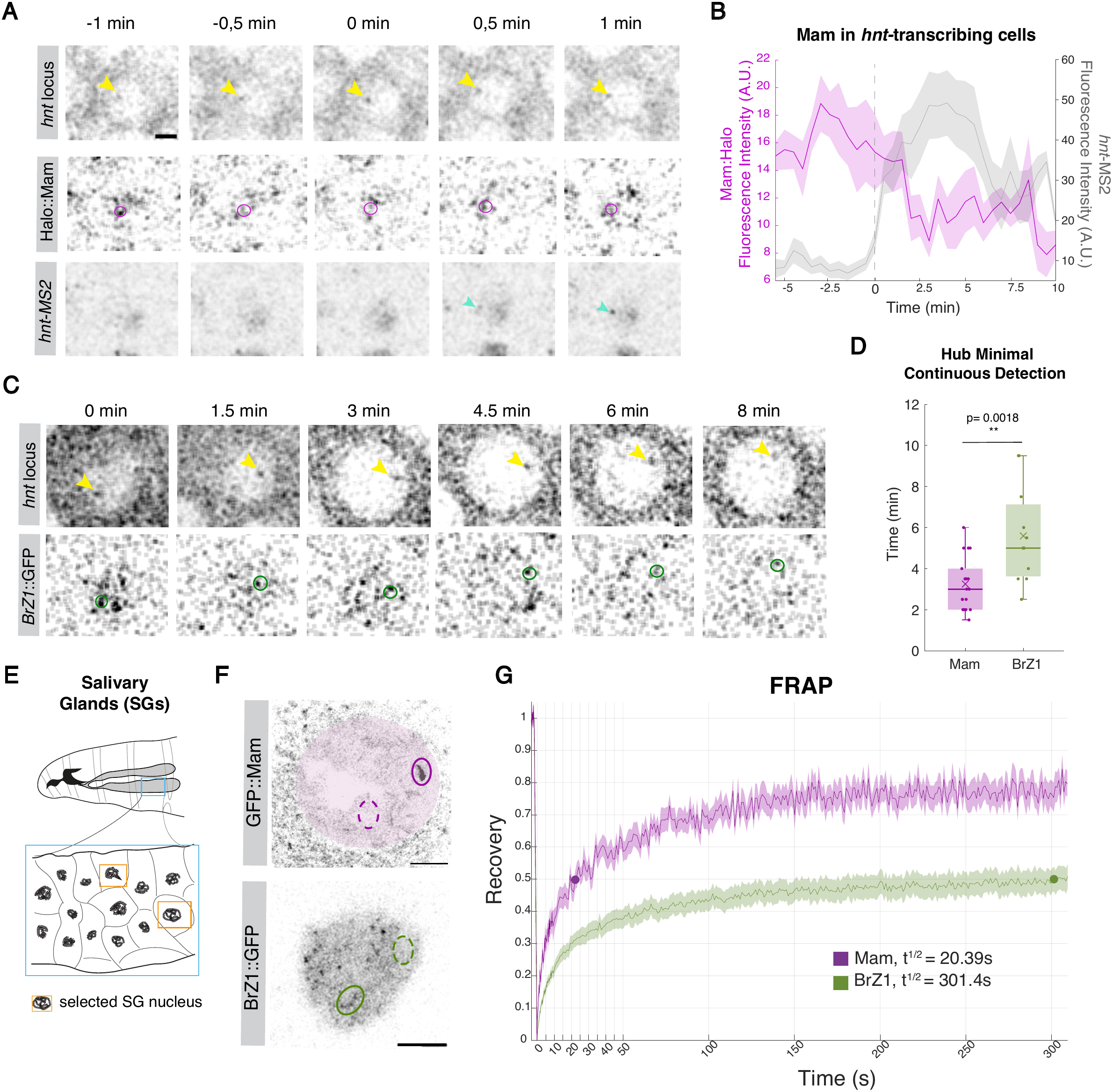
Dynamics of Mam and Broad correlate with transcription dynamics. **(A)** Confocal images from in vivo movie, tracking Halo::Mam and *hnt*-MS2/MCP::GFP in relation to *hnt* locus (yellow arrowhead). Mam enrichment (magenta circle) precedes *hnt*-MS2 transcription (cyan arrowhead). Scale bar, 2μm. Movie S3. **(B)** Average intensities of Mam (pink) and *hnt*-MS2/M-CP::GFP (grey). Intensities are time aligned respect to transcription initiation (dotted line), n = 12 nuclei from 5 e.c. **(C)** Confocal images from *in vivo* movie, tracking BrZ1::GFP (green circle) in relation to *hnt* locus (yellow arrowhead). Scale bar, 2μm. Movie S4. **(D)** Quantification of Hub Minimal Continuous Detection times for Mam (pink) and Broad (green). Boxplots indicate median, with 25–75 quartiles; error bars are SD. **(E)** Cartoon illustrating imaging approach using *Drosophila* salivary glands (SGs) with region outlined in blue amplified below. Individual nuclei (orange box) were selected for FRAP analysis. **(F)** Regions tested by FRAP for GFP::Mam and BrZ1::GFP, hub region (solid lines), control regions (dotted line). For Mam experiments, Notch-ON cells were used and hub corresponded to *E(spl)-C*. Movies S5 and S6. **(G)** Dynamics of GFP::Mam (magenta) and BrZ1::G-FP (green) in SGs nuclei analysed by FRAP. Dots indicate 50% recovery. Mam showed faster recovery than BrZ1. Legend summarizes time to 50% recovery (t1/2). Error represents the standard error of the mean (SEM).

To determine the average duration of Mam hubs at *hnt* over multiple S6 nuclei, we applied an intensity threshold to define hub detection, and calculated the minimum period over which each hub was continuously tracked (Figure 5D, Movie S4). This yielded an average duration of 3 minutes for Mam hubs, and likely included both productive and non-productive hubs. When we performed the same analysis for BrZ1::GFP at S9, we found that BrZ1 hubs were present continuously for substantially longer. The average detection period was almost twice as long as for Mam hubs and in some cases BrZ1 hub durations were as long as 10 minutes (Figure 5C,D). Notably, the difference between the Mam and BrZ1 hub durations aligns well with the difference between the Active Periods of *hnt* transcription at the two stages. This supports a direct role for hub properties in shaping the transcriptional response.

The different hub properties might reflect differing underlying dynamics of Mam and BrZ1 molecules. To investigate, we turned to the salivary gland, a tissue with large nuclei which are amenable to analysing transcription factor dynamics by Fluorescence Recovery After Photobleaching^17,50^ (FRAP) and which express both Mam and BrZ1 (Figure 5E). Under conditions of constitutively Notch activity (ectopic NΔECD) Mam was recruited to a hub at *E(spl)-C* (detectable as a band across the aligned polytene chromosomes) (Figure 5F)^17^. Nevertheless, the individual molecules remain highly dynamic as shown by the rapid timescale of recovery following photobleaching (t1/2 = 20 s) with a comparatively small immobile fraction (<20%; Figure 5G, Movie S5). In contrast, when similar bands of BrZ1 were photobleached, the rate of recovery was substantially slower than for Mam (t1/2 = 301 s) and failed to approach full recovery over the time-course of the experiment (Figure 5G, Movie S6). This suggests that >50% of the molecules were comparatively immobile over this time-period, indicating that BrZ1 complexes have a longer residence time compared to Mam complexes^51^.

Together, these results demonstrate that Mam and BrZ1 concentrate into local transcription hubs with differing durations that may be explained by their different binding properties and that correlate with the temporal profiles of the cognate *hnt* transcription at the stages when each is active.

### Mam and Broad participate in a transcription-factor handover within a hub

The two phases of *hnt* transcription can be explained by distinct regulatory inputs from Notch and BrZ1, acting on different regulatory elements. We therefore investigated the relationship between the two and whether they are required to act in an ordered sequence. To test this, we expressed the BrZ1 isoform during earlier stages and examined the consequences on *hnt* transcription by smFISH in S3 and S4 prior to the onset of Notch activation. At physiological levels, BrZ1 was insufficient to induce *hnt* expression at S3 or S4, arguing that it is insufficient to promote *hnt* transcription prior to the period of Notch activity (Figure 6A,B). However, with much higher levels of BrZ1 over-expression the restriction was overcome, and *hnt* transcription could be induced even at the early stages (Figure 6A,B). However, the Notch responsive *hnt^peb-3^* enhancer was not induced at S3,S4 under these conditions confirming that BrZ1 acts through a different regulatory element (Figure S4C). Overall the results indicate that, in normal circumstances, Notch activity is required to “prime” *hnt* so that it can respond to the low/moderate levels of BrZ1 that are present. But in situations where BrZ1 is present at high concentration, this priming is not required and BrZ1 can bypass the Notch-dependent step.

**Fig. 6.**
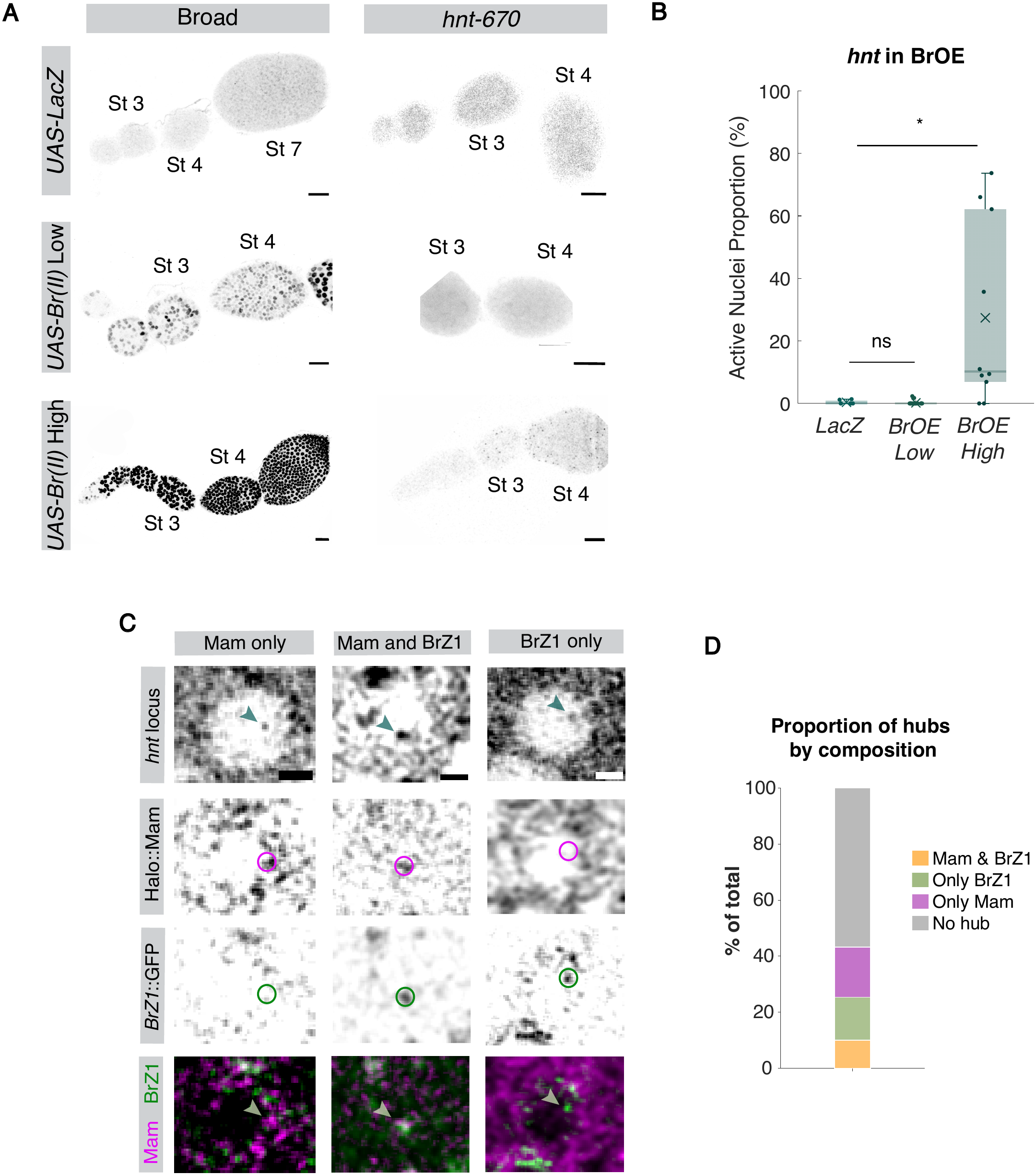
Visualizing localized TF handover at *hnt* in relation to two phase response. **(A)** Left panels: Expression of BrZ1 detected by anti-Br(core) in control (*UAS-LacZ*), low level BrZ1 (*UAS-BrZ1(II)* 6hrs) and high levels BrZ1 (*UAS-Br(II)* 24hours). Right pannels: *hnt* transcription levels detected by smFISH using *hnt*-570 probes at early stages (S3,S4) from equivalent conditions. Scale bars, 20 μm. *hnt* ATS are only present in high BrZ1 conditions. **(B)** Proportion of nuclei with *hnt* ATS under the conditions indicated, as in A, with data from S3 and S4 combined. Boxplots indicate median, with 25–75 quartiles; error bars are SD. **(C)** Confocal images, localization of Halo::Mam (magenta circle), BrZ1::GFP (green circle) with respect to *hnt* locus (arrowhead). Scale bars,2 μm. **(D)** Percentages of cells with different hubs compositions as indicated.

The sequential phases of *hnt* activation that occur normally suggest there is a handover mechanism between the two regulatory inputs. To visualise this handover we used three-colour imaging to simultaneously detect Halo::Mam, BrZ1::GFP, and the *hnt* locus. We focussed on S7 when the regulation transitions between the two and when both factors are detectably enriched at *hnt* (Figure 4F, S4A). Strikingly, three classes of hub were detected: hubs containing Mam alone, hubs containing BrZ1 and Mam together and hubs containing BrZ1 alone (Figure 6C,D). This result demonstrates that Mam and BrZ1 can coexist within a single hub, supporting a handover mechanism in which the composition of the transcription hub changes over time to confer different properties on *hnt* transcription.

## Discussion

Combinatorial TF action is a common strategy for generating accurate spatial gene expression profiles during development^52–54^. Diverse inputs must also be combined over time to confer transcriptional responses that are sustained throughout developmental processes that take place over comparatively longer periods^31^. Here, by combining analysis of transcription dynamics with live imaging of TFs at single-cell resolution in an endogenous Notch signalling context, we show that a temporal handover in TF composition within a locus specific transcription hub can explain the control of gene expression over time in a developing tissue. Furthermore, the different TF dynamics influence the properties of the hub and, in turn, those of mRNA production. Together, our findings point to TF hubs as signal integrators that coordinate transcription dynamics over time, enabling the prolonged responses that drive developmental processes.

### Role of hub kinetics in coordinating the transcriptional response

Transcription of *hnt* in the follicular epithelium comprises two dynamically distinct phases that coincide with a change in control from a Notch-responsive enhancer, *hnt^peb3^*, to one or more Notch-independent enhancers regulated by Broad. Notably, prior Notch activity is required for Broad to take over control of the *hnt* response in its late stage. The two phases are coincident with the enrichment of Mam and Broad, respectively, within a hub formed at the target locus. Indeed at the transition stage 7, we could detect recruitment of both factors into a single hub. Sequential recruitment of TFs into locus specific hubs is likely a widespread mechanism coordinating enhancer handover and integrating information over time^17,55^.

During the first phase of *hnt* regulation, the duration and levels of Mam hubs correlate with those of transcription, in agreement with our results for *E(spl)-C* genes^18^. This suggests a direct relationship between the dynamic properties of the condensate and the active periods of subsequent transcription. In support of this model, the difference in perdurance between Mam and Broad hubs recapitulated the difference between the active periods that they confer. The fact that Broad appears to have a longer residence time, based on its slower recovery in FRAP experiments, makes it likely that differential hub behaviour arises from the molecular kinetics of the TFs. In keeping with these observations, simulations have indicated that molecular binding kinetics and binding site number may be key components conferring enhancer-specific hub properties^5^. However, our finding that Mam hubs at *E(spl)-C* and *hnt* have similar characteristics, yet non-similar genome organisation (*hnt^peb-3^* is located ∼100 Kb away from the *hnt* TSS) and motif numbers, argues against the latter being the sole driver. Our observations place more emphasis on the dynamic features of the TF complexes themselves, and lead us to propose a model in which these features control hub kinetics and, in turn, the parameters of the transcriptional response.

The model that hub formation is intrinsic for Notch target-gene activation fits with other studies proposing that hubs activate transcription^24,55–58^. Our results argue for a more profound role than that implied by a recent report in which Dorsal hubs are proposed to enhance transcription efficiency without being required for activation^59^. Importantly, by tracking the target locus independently of transcription, we have measured TF enrichment before transcription (MS2/MCP signal) appears and in nuclei where no transcription is initiated, under unaltered physiological conditions^18^ . Considering two different TF complexes we find a striking relationship between the existence and perdurance of the hubs they form and the dynamics of the cognate transcription profiles. This best fits a model whereby the properties of TF hubs regulate the probability, duration and efficiency of transcription.

### TF handover mechanism orchestrates dynamic changes that sustain prolonged responses

Prolonged expression of *hnt*, lasting more than 25 hours^60^, is required for cells to transition from an immature epithelium to a mature, ovulation-competent follicle^35–37^. It depends on the activity of Broad which is itself proposed to be a target of the earlier Notch activation complex^46^ and is insufficient to activate *hnt* transcription on its own at physiological levels. This relationship implies a feed forward regulatory loop, that ensures and reinforces the handover between Notch and Broad. A similar feed forward handover relationship occurs during muscle differentiation to coordinate spatio-temporal regulation of key patterning genes^30,61^ and may thus be a widespread regulatory motif in temporal gene expression programmes. Explanations for how this sequential TF action might operate are that the Notch activator complex brings chromatin remodellers that prime the second enhancer(s) to respond to Broad^16^ or that sparse transcription during the first phase maintains a competent state, for example through nucleosome eviction^62^. Whichever the mechanism, it enables a temporally orchestrated progression in *hnt* transcription.

*hnt* transcriptional dynamics change during this process. At early stages, *hnt* transcription is stochastic, with proportions of active nuclei and active times similar to those of the *E(spl)-C* genes at equivalent stages^18^, consistent with Notch regulating this first phase. After stage 7, *hnt* transcription becomes more sustained, with a larger proportion of active cells, longer active periods and higher amplitude. This is consistent with the proposal that loci controlled by multiple enhancers can be regulated through burst duration as well as by burst frequency^6^ and with other examples where burst duration is regulated during development^7,63^. Because noise in mRNA levels increases with the variability of ON and OFF durations, tight control of burst duration is predicted to buffer stochastic variation in expression^6^. According to our model, handover from a dynamic complex to a more stable TF would lengthen burst duration, providing temporal control, and reduce transcriptional noise. It may thus be an important step in stabilising cell states.

Together, our findings point to changes in the composition of a locus-specific hub as a mechanism by which cells integrate signals over time and inform prolonged transcriptional responses. Therefore, linking hub composition and properties to the characteristics of transcription may provide a general strategy for adapting transcriptional responses in time, enabling cells to acquire and maintain their identity during differentiation.

## Materials G Methods

### Experimental animals

Species: *Drosophila melanogaster*. Flies were grown and maintained on food consisting of the following ingredients: glucose 76 g/l, cornmeal flour 69 g/l, yeast 15 g/l, agar 4.5 g/l, and methylparaben 2.5 ml/l.

### Fly stocks

Full genotypes are summarised in Table S1 and S2. For MCP/MS2 live imaging of transcription, *hnt-*MS2^41^ was combined with *hsp83-MCP::GFP* (Bloomington *Drosophila* Stock Center, BDSC #7280) to track transcription and His2Av-RFP (BDSC #23650) to label nuclei.

To image *hnt* locus a *hnt* “locus tag” strain was generated by inserting *Int1* sequence into *hnt* upstream regulatory region (Chromosome X :4613903) by CRISPR engineering^7^ (see below). *hnt* “locus tag” chromosome was combined with *UAS-ParB1-mCherry* inserted *AttP.8CFb* and *traffic-jam-Gal4* to drive ParB1 expression in the FE^64^. To analyse expression and enrichment of TFs and other proteins we used endogenously tagged fluorescent labelled proteins generated by CRISPR, Halo::Mam, GFP::Mam^17^ , Dl::mScarlet ^65^, and the PBac BrZ1::GFP (BDSC #50754). For triple labelling experiments *hsp83-MCP::GFP* (BDSC #7280) was recombined with traffic-jam(tj)-Gal4 to generate a strain containing Halo::Mam, *hnt* “locus tag”UAS*-ParB1-mCherry* and *hnt*-MS2. FRAP experiments in the larval salivary glands were performed by crossing *1151-Gal4*, *to* GFP::Mam, *UAS-NΔECD* or to BrZ1::GFP.

To manipulate protein functions, RNAi lines were as listed in Table S1 and included *UAS-Hairless-RNAi* (BDSC #27315), *UAS-Broad-RNAi* (BDSC #51189) and *UAS-mirr-RNAi* (BDSC #65877). Controls as appropriate for third chromosome was *UAS-white-RNAi* (III). All of these were by combined to tj-Gal4::tub-Gal80^ts^. For Broad overexpression, UAS-BrZ1(II) or UAS-LacZ as controls were used in combination to tj-Gal4::tub-Gal80^ts^. Crosses were maintained at 18°C (non-permissive temperature due to Gal80 repression) or moved to 29°C (permissive temperature due to Gal80 repressor downregulation) for 72 hours for RNAi expression, 6 hours or 24 hours for Low or High Broad overexpression, respectively.

## Method details

### Generation of tagged *hnt* locus

To label *hnt (peb/CG12212)* locus, we used the Par/Int system in which CRISPR/cas9 genome editing was used to introduce *Int1* sequence into non-conserved fragment of *hnt* upstream regulatory region (Chromosome X :4613903). Briefly, one gRNA sequence(s) was cloned into U6 promoter plasmid. Cassette *int1-PBacDsRed*, which contains int1 and a *3xP3-DsRed* flanked by PiggyBac terminal repeats, and two homology arms were cloned into pUC57-Kan as donor template for homologous repair. *peb/CG12212*-targeting gRNAs and hs-Cas9 were supplied in DNA plasmids, together with donor plasmid and microinjected into embryos of control strain *LWG228 w[1118]*. Transformants were selected based on the expression of 3xPax3-dsRED and confirmed by PCR and sequencing.

CRISPR Target Site [PAM]: TGTCGTCGCAGTGCCATGAA[TGG]

Guide RNA Primers: Sense oligo 5’-CTTCGTGTCGTCGCAGTGCCATGAA

Upstream Homology Arm: 974bp, from X:4612928 to 4613901

Forward Oligo5’-AACTGATTCATGGCCACACG

Reverse Oligo5’-TGGCACTGCGACGACACC

Downstream Homology Arm: 1,049bp, from X:4613906 to 4614954.

Forward Oligo: 5’-TGGTTGGGTCATAAATGCAA

Reverse Oligo: 5’-GCCAGCGAATTTGTTATTG

All this protocol was performed by WellGenetics Inc. using modified methods of Kondo and Ueda (2013).

### Single Molecule in Situ Hybridisation

Custom smFISH probe sets for *hnt, E(spl)m7-HLH*, *E(spl*)*mβ*-HLH and *E(spl)m3-HLH* genes were designed with Stellaris Probe Designer (Bioscience Technologies). Ovaries were dissected in non-supplemented Schneider’s medium and fixed for 30 min in 3.7% formaldehyde:PBS at room temperature. Ovaries were washed three times in PBST (PBS +0.1% Triton X-100) and transfered to Methanol using increasing concentrations with 5 min incubations in PBST:MeOH 7:3, PBST:MeOH1:1, PBST:MeOH 3:7 on a nutating mixer before transferring to 100% MeOH. After 10 mins in 100% MeOH, tissues were rehydrated with the same solutions in the reverse order (3:7 PBST:MeOH, 1:1 PBST:MeOH, 7:3 PBST:MeOH). After rinsing with PBST, ovaries were prehybridized with WB-A (Biosearch Technologies SMF-HB1-10), for 10 min and then transferred to hybridisation mixture prepared by adding probes to 100 µl of RNA FISH Hybridization Buffer (Biosearch Technologies SMF-WB1-60) at 1:100 %v/v from 12.5 µM stock. They were then incubated at 37°C for 16-18 hours in the dark, washed twice for 30 minutes with pre-warmed WB-A, rinsed with PBST and then mounted using Vectashield with DAPI.

smFISH samples were imaged using a Leica SP8 microscope equipped with different laser lines (405, 458, 488, 496, 514, 561, 594 and 633) and a 40x apochromatic 1.3 oil immersion objective and one hybrid GaAsP detector was used. Individual egg chambers were imaged with a 4x zoom, 1024x1024 pixel resolution, pinhole set to 1-Airy, three line-averages, a 12-bit depth, and 400 Hz scanning speed. Z-stacks were 0.5 µm width and a total of around 30 stacks.

### Tissue preparation and live imaging

Ovaries were dissected and mounted in Schneider’s medium (Sigma-Aldrich/Merck, S0146) supplemented with 15% (v/v) fetal bovine serum (Sigma-Aldrich, F2442), 0.6% (v/v) streptomycin/penicillin antibiotic mix (Invitrogen, 15140-122), and insulin (0.20 mg/ml; Sigma-Aldrich, 15500). For live imaging, ovaries were mounted into 35-mm poly-d-lysine–coated glass-bottom dishes (Mattek, P35GC-1.5-10-C). Live confocal fluorescence imaging of ovaries was performed on a Leica SP8 microscope equipped with seven laser lines (405, 458, 488, 496, 514, 561, and 633) and using a 40× apochromatic 1.3 oil immersion objective and two hybrid GaAsP detectors.

For Mam and Broad enrichment assays, individual egg chambers were imaged with a 4x zoom, 512x512 pixel resolution, pinhole set to 1.5-Airy, three line-averages, a 12-bit depth, and 400 Hz scanning speed. Z-stacks were 0.5 µm wide and a total of circa 30 stacks were acquired to cover most of the epithelium at the imaged surface of the egg chamber. For MS2/MCP movies, individual egg chambers were imaged at 3.5 zoom, 512x512 pixel resolution, pinhole set to 3.5-Airy, 8-bit depth and 400Hz scanning speed. Z-stacks of 0.5 µm width for 30 stacks were acquired. Time frame was 20 seconds. The same settings for MCP-GFP detection: 29.36mW 488nm argon laser detected with a hybrid detector.

For Mam dynamics together with MS2/MCP, egg chambers were imaged using at 4.5 zoom, 512x512 pixel resolution, pinhole set to 1.5-Airy, line average of three, 12-bit depth and 700Hz scanning speed. Z-stacks were 0.5 µm wide and circa 13 stacks were acquired. Time frame was 30 seconds. For Broad at *hnt* locus movies, pinhole was closed to 1 Airy and line average reduced to 2 to avoid bleaching.

For *Broad-RNAi* validation experiments, individual egg chambers were imaged with a 3.50× zoom, 512x512 pixel resolution and a 12-bit depth. The microscope settings cover also three line averages, a 400-Hz scanning speed and a pinhole value of 1.00 AU. Z-Stacks were 0.5 μm wide, and a total of around 30 stacks were acquired.

### Immunofluorescence

Ovaries were dissected in no-supplemented Schneider’s media and fixed in 4% paraformaldehyde (PFA) for 20 minutes, washed three times with PBT-Triton 0,1% (PBS + 0.1% Triton 20), permeabilized in PBT-Triton 1% for 1 hour and then blocked in PBT10 (PBT + 10% BSA), rollling for 1 hour. Incubation with the primary antibody was done in PBT1 (PBT + 1% BSA) overnight at room temperature, ovaries were then washed four times for 30 minutes each with PBT1 and then incubated with the secondary antibody (1:200, Jackson immunoresearch) in PBT0.1 (PBT + 0.1% BSA) for 2 hours at room temperature. After washing three times for 10 minutes each in PBT, the ovaries were mounted in Vectashield with DAPI. For visualization of *hnt^peb-3^*:GFP reporter Rabbit anti-GFP (1:500) (Thermo Fisher **#**A-11122 ) primary and anti-Rabbit FITC (1:200, Jackson ImmunoResearch Laboratories, Inc and Sigma) secondary antibodies were used. For Broad levels detection experiments Broad core (1:25) (25E9.D7, DSHB) and anti-Mouse FITC (1:200, Jackson ImmunoResearch Laboratories) were employed. Image acquisition was performed in a Leica SP8 microscope equipped with four laser lines (405, 488, 552, and 638) and one hybrid GaAsP detector. Image acquisition was similar to live imaging but with a with an optimised Z step size, different zooms and a 512x512 px resolution.

### Image analysis pipeline

For smFISH images, nuclei were segmented in 2D using Weka Segmentation (Fiji plugin) to detect DAPI staining and masks were generated using watershed after transformation to a binary image and their number indicated the total number of nuclei analyzed per egg chamber. Masks were applied to the fluorescent probe channels to eliminate signal coming from the cytoplasm and a minimal threshold was used to identify the prominent ATS. The Fiji plugin Analyze Particles was used to count total number of ATS in an image and totals were divided by the number of nuclear masks and represented as proportion of Active Nuclei.

To quantify the fold-change difference of two conditions the following protocol was applied. For each developmental stage, the difference between conditions was quantified as the log2 fold change of group means. Because the conditions consisted of independent (unpaired) samples, uncertainty was estimated with a nonparametric percentile bootstrap. For each stage, 5,000 bootstrap resamples were generated by sampling with replacement from the observed values of each condition independently, preserving the original group sizes. The log2 fold change of means was recalculated for every resample, and the 95% confidence interval was defined as the 2.5th and 97.5th percentiles of the resulting bootstrap distribution. The plotted point estimate is the log2 fold change calculated from the original data rather than the bootstrap mean. A log2 fold change of 0 indicates no difference between conditions.

For MS2/MCP movies, the His2Av-RFP signal was used to segment and track the nuclei in 2D. Each movie was segmented using Weka Segmentation (Fiji plugin) and watershed after transformation to a binary image to obtain masks. Tracking of the nuclei was performed using Trackmate (Fiji plugin) detecting the nuclei as masks and allowing to track a maximum distance of 3 µm, a maximum gap distance allowed of 3 µm and maximum frame gap of 2 frames. A custom MATLAB script (MATLAB R2021b, Mathworks), adapted from previous work from the lab^7^, was used to detect and track the transcription spots based on the 3D Gaussian method developed in ^40^. The tracked transcription spots were overlapped with the ROIS (masks) of the tracked nuclei, and spots lying outside nuclei were removed from the analysis.

For *Broad-*RNAi validation efficiency experiment, three egg chambers were considered for each developmental stage. Three representative frames were selected from a series of around 30 stacks and for each plane were considered 10 nuclei centrally located using the Oval Selection in order to obtain approximately 90 nuclei per stage. The different values of Raw Integrated Density were normalized to the nuclei area to determine the mean fluorescence intensity, from which was substracted the background fluorescence measured in 9 nurse cells from each developmental stage.

For Broad knockdown in *hnt^peb-3^*:GFP reporter background, a square of identical size was drawn at the centre of each egg chamber images, which were previously converted to maximum-intensity projections in Fiji. From the ROI obtained for each image, the different values of Raw Integrated Density were normalized to the area to determine the mean fluorescence intensity.

To measure enrichments, images were analysed using MATLAB by importing Leica Images with BioFormat package (MATLAB R2021b, MathWorks and openmicroscopy.org). Using a custom MATLAB app, a squared region of interest (ROI), 30 × 30 pixels centred on the *hnt* locus was selected (adapted from ^17^). Fluorescence levels of Halo::Mam and BrZ1::GFP were then measured within this ROI, averaged per experimental condition and normalized with an average value, calculated from ROIs placed at random in the same set of nuclei and represented as mountain plots. For quantification of Mam enrichment together with MS2 and BrZ1 movies, *hnt* was tracked with Trackmate and the resulting ROI was assigned to the Mam and MS2 channels and fluorescent intensities obtained using a custom MATLAB script. Both Mam and MS2 signals were normalized by subtracting the background level (measured as an average of 3 regions with no hub or MS2) and median filtered to smooth the traces. To determine the hub minimal continuous detection for both Mam and BrZ1, individual thresholds were applied to every cell depending on background levels in each movie and a maximum of two frames gap were allowed to count as a single trace.

For quantification of proportions of hubs, thresholds for each protein were applied to average signal of a reduced area (10x10 pixels) centered at the *hnt* locus. The thresholds were calculated using the limit value for quantiles showing discrete enrichment at the center of the region.

### FRAP experiments

FRAP experiments were performed on Leica SP8 microscope equipped with seven laser lines (405, 458, 488, 496, 514, 561, and 633) and bleaching and scanning speed were optimized: pinhole set to 3.5-Airy, speed was increased to 700 Hz, no line averaging was used, and Leica FRAP booster was activated. Point bleaching was employed. Images before (10 images) and after bleaching were acquired every 0.32 s. After 50 images post bleaching, frame gap was increased to 2s to minimise bleaching. Bleached regions were selected to show high Mam or BrZ1 signal including regions with bands. Control regions with low enrichment and located distantly from the bleached zone were used to normalise for acquisiton photobleaching, as described^16^ by applying the following formula:

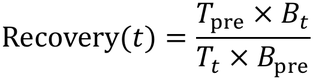

where:

- *B*_t_: mean fluorescence intensity of the bleached region throughout the experiment
- *B*_pre_: mean fluorescence intensity of the bleached region before bleaching
- *T*_t_: mean fluorescence intensity of the whole nucleus throughout the experiment
- *T*_pre_: mean fluorescence intensity of the whole nucleus before bleaching

### MS2 Data Processing and parameters calculation

MS2/MCP transcription foci with a minimum of 5 frames were analysed allowing for a maximum gap of 2 frames. This generated an initial distribution of active (nuclei transcribing for a minimum of 5 frames) and inactive nuclei. Kymographs only displaying active nuclei over time were produced for each movie. Only traces that started after the nuclei had been segmented and finished before their track ends were kept for analysis to avoid false negatives arising from nuclei moving out of the field of view. During the Active periods, short gaps (less than 5 frames) were included in the trace, and their mean intensity was interpolated using the mean intensity value of the surrounding frames. The active nuclei proportion was calculated as the percentage of nuclei that have transcription puncta per frame. The amplitude of MS2 transcription was defined by aligning the onset of the Active periods and to account for proportion of active nuclei as well, values of 0 were assigned when the trace ended.

Active period durations were calculated using the same procedure, except short gaps were permitted if they lasted fewer than 21 frames (approximately 7 minutes). Only Active Periods between two Inactive Periods were counted. In a similar way, inactive periods between two active ones were quantified.

### MS2 signal calibration

To calibrate the signal from MS2 foci from *in vivo* movies and obtain an average number of mRNA molecules present in them, we calculated the fluorescence of a single mRNA molecule, according to established method^18,25,66^.

### Modelling

Signal deconvolution and kinetic model inference were performed using the BurstDECONV framework^26^. Calibration, software application and modifications as described in (^18^).

### Statistical Analysis

Two-tailed *t*-test was applied in cases with n > 30 for both conditions tested. Otherwise, normality was checked via a Shapiro–Wilk test. When one or more of the samples were not normal, a Wilcoxon rank sum test was applied. In all cases, significance was presented as: *p<0.05, **p<0.01, ***p<0.001, and ****p<0.0001, and p-values are provided in Figure legends.

## Data and code availability

https://doi.org/10.5281/zenodo.24133649

## Acknowledgements

We thank our colleagues and members of our group for discussions and helpful suggestions. We are grateful to Javier De Haro Arbona for his help with FRAP experiments, to Kat Millen and Emilia Campanella for assistance and technical resources, to Maria Jesús Gómez Lamarca and Violeta García Hernández for reading and commenting on the manuscript and to Cambridge Advanced Imaging Centre for microscopy support.

## Author contributions

Conceptualization: CSM, CR, SJB

Methodology: CSM, CR, SJB

Software: CSM, CR

Validation: CSM, CR

Formal Analysis: CSM, CR

Investigation: CSM, AM

Resources: CSM

Visualization: CSM, CR, AM, SJB

Supervision: SJB, CSM

Writing—original draft: CSM, CR, SJB

Writing—review & editing: CSM, CR, SJB

Project Administration: SJB

Funding acquisition: CSM, SJB

## Data and materials availability

All data needed to evaluate the conclusions in the paper are present in the paper and/or the Supplementary Materials, raw images and quantification are deposited in FigShare: https://figshare.com/projects/Transcription_factor_handover_reshapes_hub_dynamics_to_coordinate_temporal_progression_of_Notch_response/284542

Materials generated in this study are available on application to the corresponding author. Code are available here https://doi.org/10.5281/zenodo.24133649.

## Supplementary Figures

Figure S1. Related to Figure 1. Notch activity ends at stage 8 whereas *hnt* transcription persists.

Figure S2. Related to Figure 2. *hnt* transcription active periods in stage 6 are similar to those in *E(spl)-C* genes and change in stage 9.

Figure S3. Related to Figure 3. Probing potential regulators for different *hnt* transcription phases.

Figure S4. Related to Figure 4,5 and 6. Percentage of cells with Mam hub at *hnt* locus and cross-correlation of Mam hub with *hnt*-MS2/MCP::GFP. Levels of *hnt^peb-3^-GFP* in Broad overexpression.

## Supplementary Tables

Table S1: Details of *Drosophila* lines used.

Table S2: Detailed genotypes for each Figure.

Table S3: “n” numbers and *p*-values from statistical tests for each Figure.

## Supplementary Movies

Movie S1: Live imaging of *hnt-*MS2/MCP::GFP transcription at St 6 egg chamber.

Movie S2: Live imaging of *hnt-*MS2/MCP::GFP transcription at St 9 egg chamber.

Movie S3: 3 colour live imaging of *hnt* locus (yellow), Mam::Halo (magenta), *hnt*-MS2/MCP::GFP transcription (orange), stage 6.

Movie S4: Live imaging of BrZ1:GFP (green) at *hnt* locus (yellow) at stage 9.

Movie S5: Live imaging of FRAP of Mam::GFP at salivary gland nuclei.

Movie S6: Live imaging of FRAP of BrZ1::GFP at salivary gland nucle

## Supplementary Figure Legends

**Fig. S1. Notch activity ends at stage 8 whereas *hnt* transcription persists.**

(A) *hnt* activty in relation to *E(spl)-C*, measured by proportion of nuclei with *hnt* ATS versus the mean proportion of nuclei with *E(spl)-C* gene ATS at the stages indicated. The shaded area represents a confidence interval (CI) of 95% generated from data in Figure 1C. (B) Confocal images of smFISH with *E(spl)m3-570* in control (*LacZ* expression) versus NΔECD expression at stage 9. Masks from nuclei segmentation are superposed on the images. Large increase in ATS detected with ectopic Notch activation. Scale bars, 5 μm. (C) Confocal images of smFISH with *E(spl)m3-570* in control (*wRi*) vs Hairless-RNAi (*HRi*) conditions: at stage 6, a reduced proportion of nuclei have ATS (arrowheads), at stage 9 a similar minimal number of nuclei retain ATS in *Hairless-*RNAi conditions. Scale bars, 5 μm. (D) Fold change analysis of *E(spl)m3* activation in control (*wRi*) versus *HRi*, measured by proportion of nuclei with *E(spl)m3* ATS in the two conditions. Shaded area represents a confidence interval (CI) of 95%. Data from Figure 1E. (E) Confocal images of smFISH with *hnt-C70* in control (*wRi*) vs Hairless-RNAi (*HRi*) conditions: at stage 6, an increased proportion have ATS (arrowheads), at stage 9 proportion of nuclei with ATS remains unchanged. Scale bars, 5 μm. (F) Fold change analysis of *hnt* ATS in control (*wRi*). versus *HRi*, measured by proportion of nuclei with ATS in each condition. Shaded area represents a confidence interval (CI) of 95%. Data from Figure 1F.

**Fig. S2. *hnt* transcription active periods at stage 6 are similar to those of *E(spl)-C* genes and change at stage G.**

(A) Violin plot represents active period durations in transcription traces from *E(spl)-m7* (purple), *E(spl)-mβ (*lilac) and *hnt* (turquoise), all at stage 6. (B) Example trace from *hnt* transcription at stage 9 calibrated by number of molecules and their corresponding Pol II initiation events calculated by BurstDECONV. (C) Table indicating absolute values from the different rates present in a 3-state promoter model of *hnt* at stage 6 and 9. Substantial differences are depicted in bold.

**Fig. S3. Probing potential regulators for different *hnt* transcription phases.**

(A) Confocal images from an ovarioles nuclei detected with DAPI and corresponding expression of *hnt^peb-1^*::GFP and *hnt^peb-2^*::GFP. Neither have detectable expression in the FE at any stage. Scale bars, 50 μm. (B) *Br*RNAi depletes BrZ1::GFP protein levels confirming efficacy; normalized intensity of BrZ1::GFP levels in *Br*RNAi versus *wRNAi* conditions. n= from at least 3 e.c. (C) Proportion of nuclei with *E(spl)-m7 ATS* detected by smFISH in control *(wRi*) and *Br-RNAi* (*BrRi*) conditions through stages 6, 7 and 9. (n=4-7 e.c.). No change is detected with Broad depletion. (D) Proportion of nuclei with *hnt ATS* detected by smFISH in control *(wRi*) and *mirr-RNAi* (*mirrRi*) conditions through stages 6, 7 and 9. Mirror depletion has no significant effect. n=11-18 e.c.

**Fig. S4. Percentage of cells with Mam hub at *hnt* locus and cross-correlation of Mam hub with *hnt*-MS2/MCP::GFP. Levels of *hnt^peb-3^-GFP* in Broad overexpression.**

(A) Percentages of cells with Mam hub located at the *hnt* locus during the indicated stages. n=110, 130, 85, 113, 111,100 from at least 3 e.c. (B) Cross-correlation analysis of the relationship between *hnt* transcription profiles and Mam enrichment. Mean correlation R2 values (top panels) and heatmaps (bottom panels) comparing the Mam profiles at different time lags with *hnt-*MS2/MCP::GFP intensity. (C) Expression of *hnt^peb-3^*-GFP in *Broad overexpression* (High *UAS-Br(II)*) conditions.

