## Supplementary Material for "Transcription factor handover reshapes hub dynamics to coordinate temporal progression of Notch response"

#### Santa Cruz Mateos: Supplementary Figures, Tables and movies

##### Supplementary Figures

Figure S1. Related to Figure 1. Notch activity ends at stage 8 whereas *hnt* transcription persists.

Figure S2. Related to Figure 2. *hnt* transcription active periods in stage 6 are similar to those in *E(spl)-C* genes and change in stage 9.

Figure S3. Related to Figure 3. Probing potential regulators for different *hnt* transcription phases.

Figure S4. Related to Figure 4,5 and 6. Percentage of cells with Mam hub at *hnt* locus and cross-correlation of Mam hub with *hnt*-MS2/MCP::GFP. Levels of $hnt^{peb\text{-}3}$*-GFP* in Broad overexpression.

##### Supplementary Tables

Table S1: Details of *Drosophila* lines used.

Table S2: Detailed genotypes for each Figure.

Table S3: “n” numbers and *p*-values from statistical tests for each Figure.

##### Supplementary Movies

Movie S1: Live imaging of *hnt*-MS2/MCP::GFP transcription at St 6 egg chamber.

Movie S2: Live imaging of *hnt*-MS2/MCP::GFP transcription at St 9 egg chamber.

Movie S3: 3 colour live imaging of *hnt* locus (yellow), Mam::Halo (magenta), *hnt*-MS2/MCP::GFP transcription (orange), stage 6.

Movie S4: Live imaging of BrZ1:GFP (green) at *hnt* locus (yellow) at stage 9.

Movie S5: Live imaging of FRAP of Mam::GFP at salivary gland nuclei.

Movie S6: Live imaging of FRAP of BrZ1::GFP at salivary gland nucle

### Figure S1

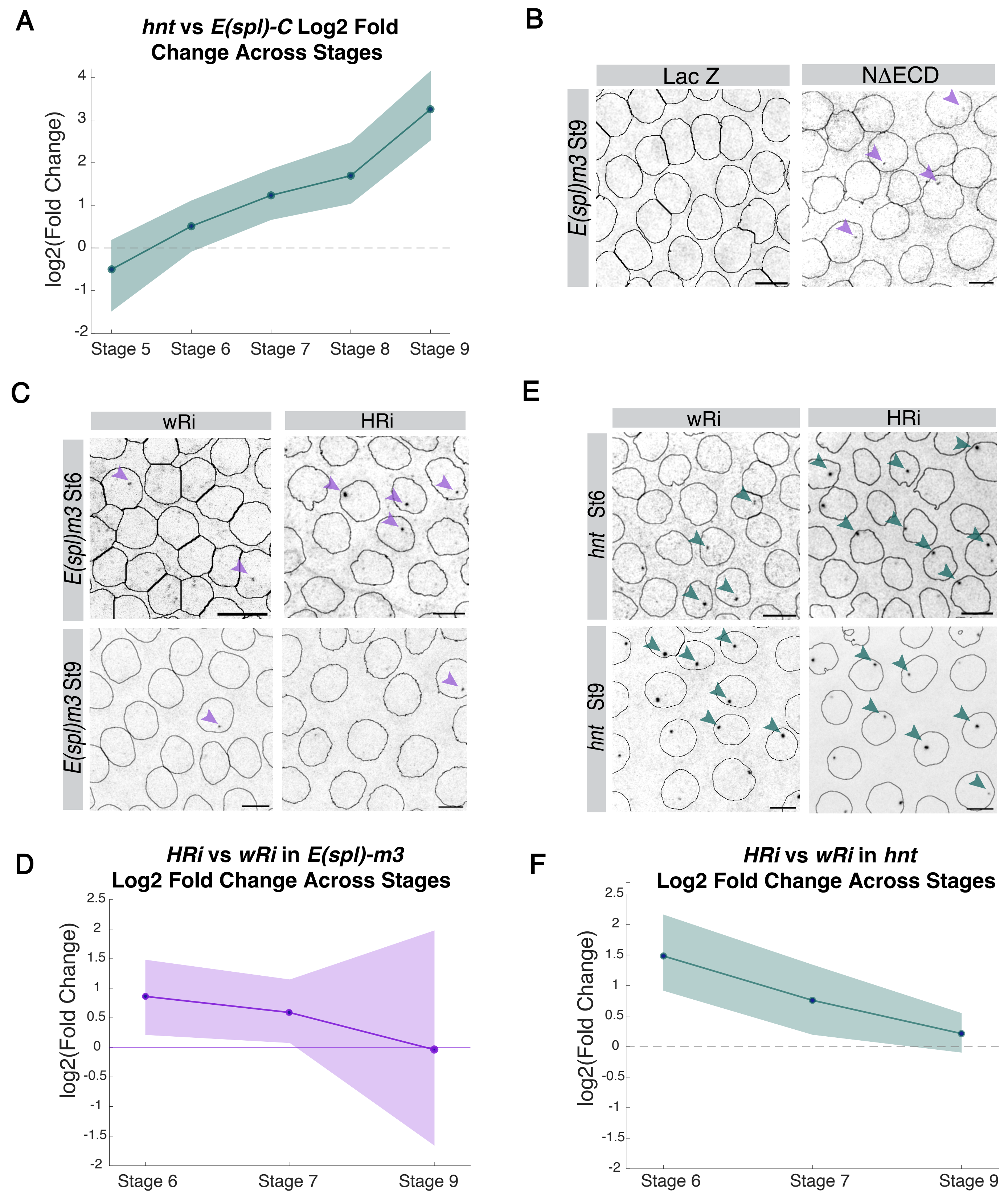


**Fig. S1. Notch activity ends at stage 8 whereas *hnt* transcription persists.**

**(A)** *hnt* activty in relation to *E(spl)-C*, measured by proportion of nuclei with *hnt* ATS versus the mean proportion of nuclei with *E(spl)-C* gene ATS at the stages indicated. The shaded area represents a confidence interval (CI) of 95% generated from data in Figure 1C. **(B)** Confocal images of smFISH with *E(spl)m3*-570 in control (*LacZ* expression) versus NΔECD expression conditions at stage 9. Masks from nuclei segmentation are superposed on the images. Large increase in ATS detected with ectopic Notch activation. Scale bars, 5 µm. **(C)** Confocal images of smFISH with *E(spl)m3*-570 in control (*wRi*) vs *Hairless-RNAi* (*HRi*) conditions. At stage 6, a reduced proportion of nuclei have an ATS (arrowheads), at stage 9 a similar minimal number of nuclei retain ATS in *Hairless-RNAi* conditions. Scale bars, 5 µm. **(D)** Fold change analysis of *E(spl)m3* activation in control (*wRi*) versus *HRi*, measured by proportion of nuclei with *E(spl)m3* ATS in the two conditions. Shaded area represents a confidence interval (CI) of 95%. Data from Figure 1E. **(E)** Confocal images of smFISH with *hnt*-670 in control (*wRi*) vs *Hairless-RNAi* (*HRi*) conditions: at stage 6, an increased proportion have ATS (arrowheads), at stage 9 proportion of nuclei with ATS remains unchanged. Scale bars, 5 µm. **(F)** Fold change analysis of *hnt* ATS in control (*wRi*). versus *HRi*, measured by proportion of nuclei with ATS in each condition. Shaded area represents a confidence interval (CI) of 95%. Data from Figure 1F.

### Figure S2

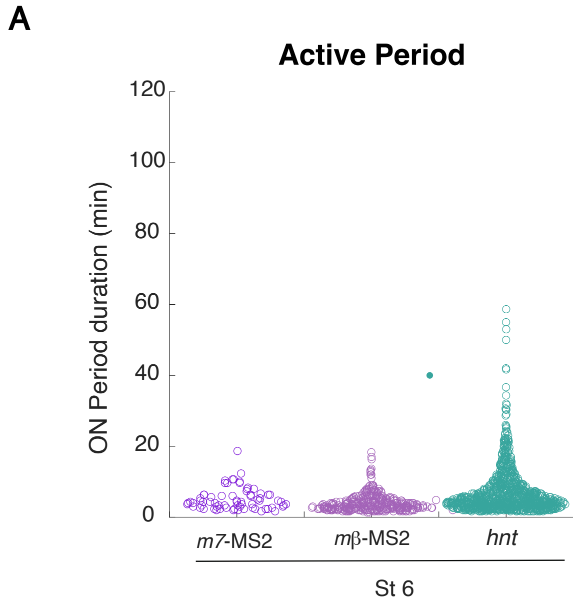


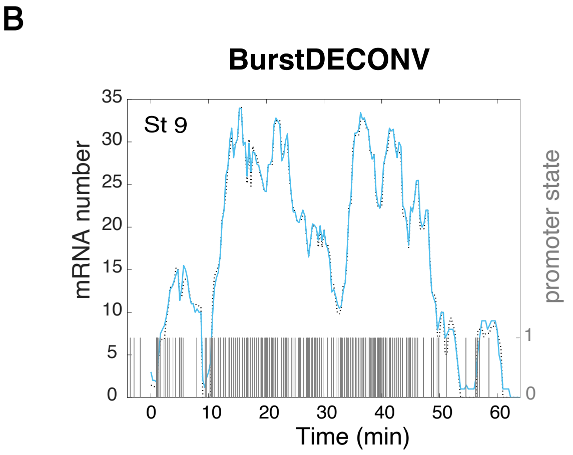


**C**

| | $k1^{+}$ | $k1^{-}$ | $k2^{+}$ | $k2^{-}$ | $k_{ini}$ |
|---|---|---|---|---|---|
| *hnt* St 6 | **2.29e-4** | 2.80e-3 | 1.62e-2 | **1.82e-2** | 5.94e-2 |
| *hnt* St 9 | **1.35e-3** | 1.11e-3 | 8.06e-3 | **5.02e-3** | 7.87e-2 |

**Fig. S2. hnt transcription active periods at stage 6 are similar to those of *E(spl)-C* genes profiles and change at stage 9.**

**(A)** Violin plot represents active period durations in transcription traces from *E(spl)-m7* (purple), E(spl)-mβ(lilac) and *hnt* (turquoise), all at stage 6. **(B)** Example trace from *hnt* transcription at stage 9 calibrated by number of molecules and their corresponding Pol II initiation events calculated by BurstDECONV. **(C)** Table indicating absolute values from the different rates present in a 3-state promoter model of *hnt* at stage 6 and 9. Differences are depicted in bold.

### Figure S3

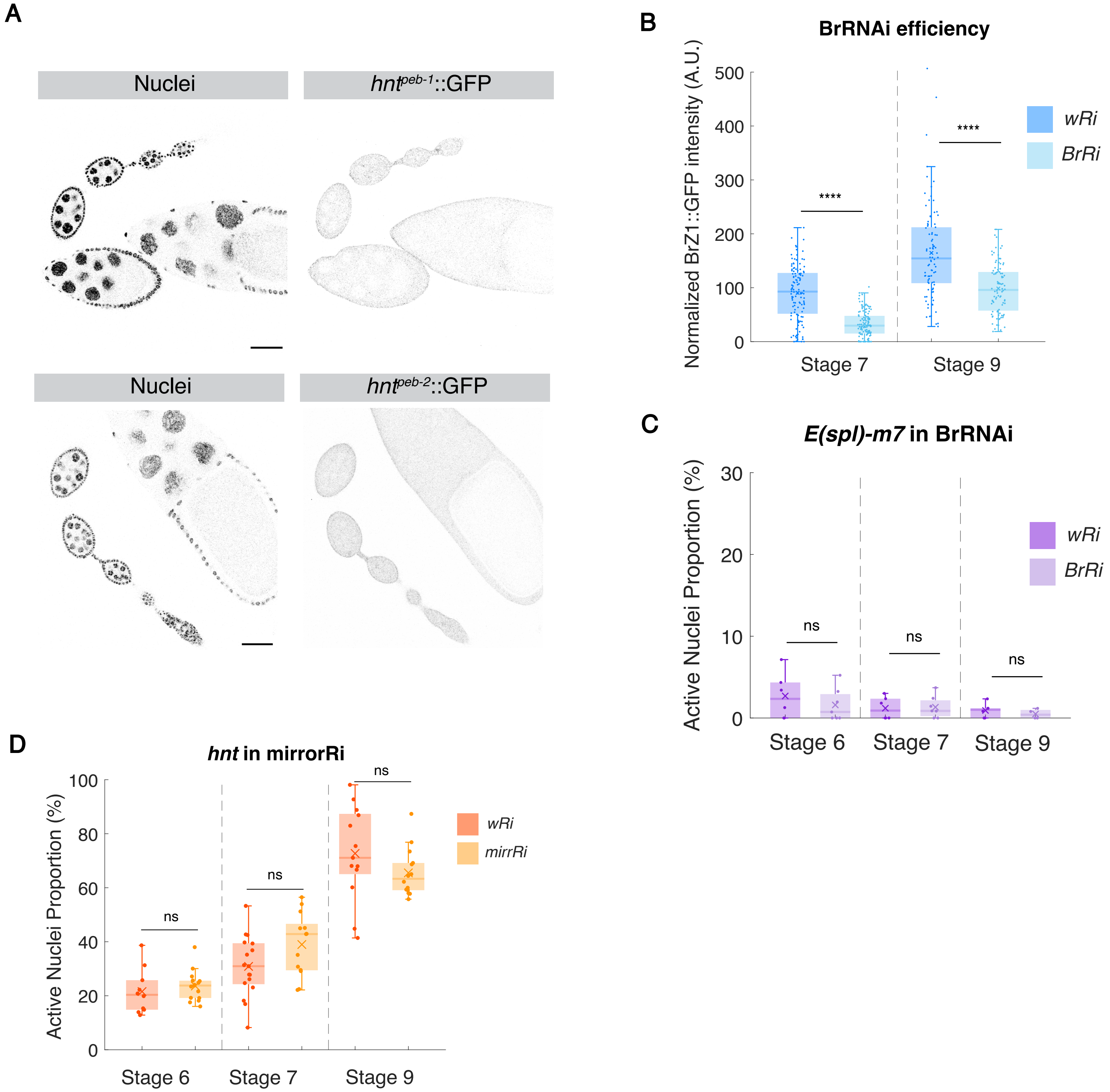


**Fig. S3. Probing potential regulators for different hnt transcription phases.**

**(A)** Confocal images from an ovarioles nuclei detected with DAPI and corresponding expression of *$hnt^{peb\text{-}1}$::GFP* and *$hnt^{peb\text{-}2}$::GFP*. Neither have detectable expression in the FE at any stage. Scale bars, 50 µm. **(B)** *BrRNAi* depletes BrZ1::GFP protein levels confirming efficacy; normalized intensity of BrZ1::GFP in *BrRi* versus wRi conditions. n= from at least 3 e.c. **(C)** Proportion of nuclei with *E(spl)-m7* ATS detected by smFISH in control (*wRi*) and *Broad-RNAi* (*BrRi*) conditions through stages 6, 7 and 9. (n=4-7 e.c.). No change is detected sith Broad depletion. **(D)** Proportion of nuclei with *hnt ATS* detected by smFISH in control (*wRi*) and *Mirror-RNAi* (*mirrRi)* conditions through stages 6, 7 and 9. Mirror depletion has no significant effect. n=11-18 e.c.

### Figure S4

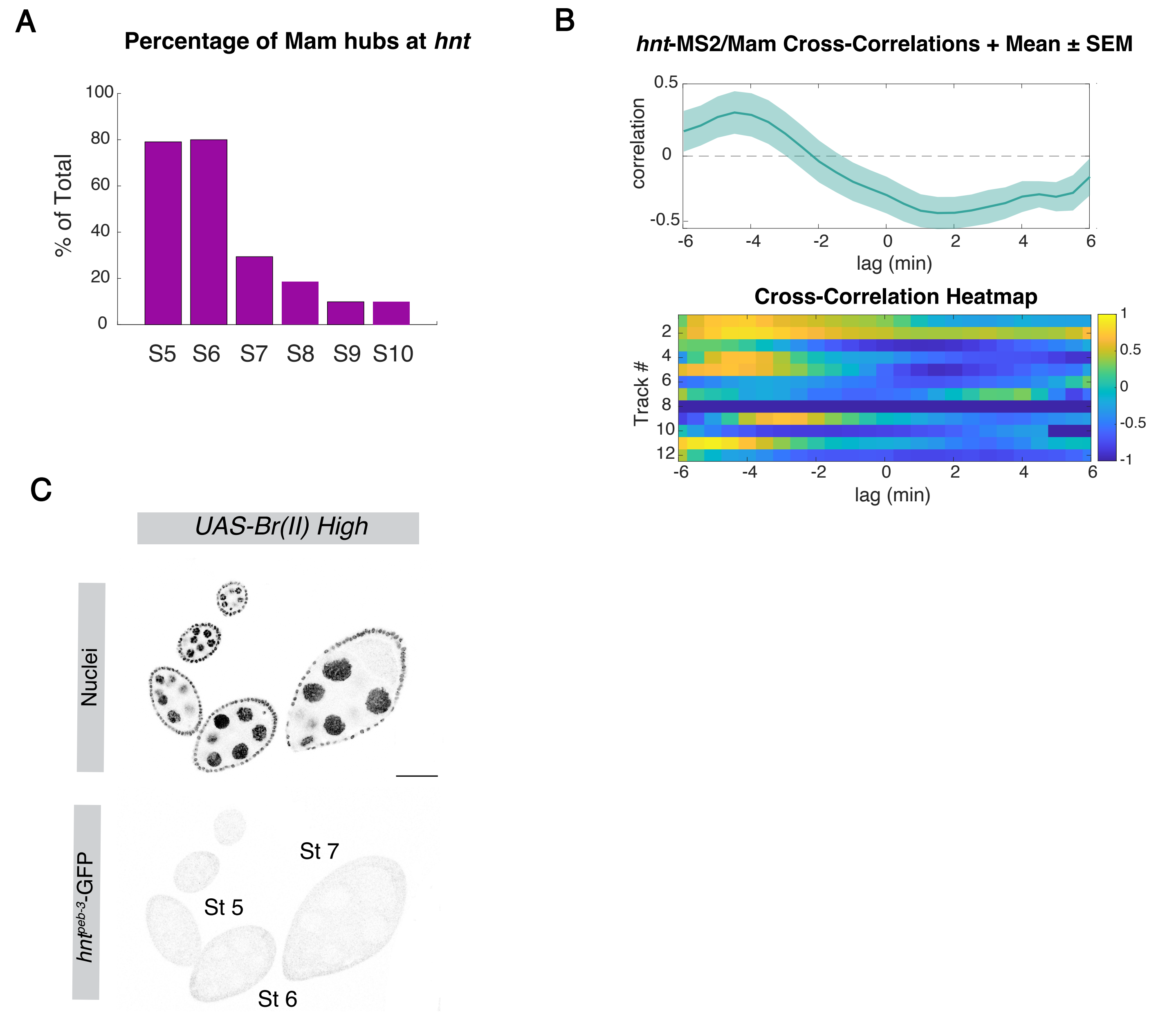


**Fig. S4. Percentage of cells with Mam hub at *hnt* locus and cross-correlation of Mam hub with *hnt*-MS2/MCP::GFP. Levels of *hnt*$^{peb\text{-}3}$-GFP in Broad overexpression.**

**(A)** Percentages of cells with Mam hub located at the *hnt* locus during the indicated stages. n=110, 130, 85, 113, 111,100 from at least 3 e.c. **(B)** Cross-correlation analysis of the relationship between *hnt* transcription profiles and Mam enrichment. Mean correlation R2 values (top panels) and heat-maps (bottom panels) comparing the Mam profiles at different time lags with *hnt*-MS2/MCP::GFP intensity.**(C)** Expression of *hnt*$^{peb\text{-}3}$-GFP in Broad overexpression (High *UAS-Br(II)*) conditions.

**Supplementary Table S1: Details of *Drosophila* lines used.**

| Line / Element | Genotype (FlyBase Nomenclature) | Source |
|---|---|---|
| *white [1118]* | *w[1118]* | |
| *tj-GAL4::tub-Gal80$^{ts}$* | *w[*], P{w[+]=GawB}Tj , P{w[+]=tubP-GAL80[ts]}* | St Johnston lab |
| *UAS-NΔECD* | *P{w[+]=UAS-NotchΔECD,* mCherry *}* | Rebay et al., 1993; Fortini et al., 1993 |
| *UAS-LacZ* | *P{w[+]=UAS-nls-LacZ}* | Brand Lab |
| *UAS-Hairless-RNAi* | *y1 v1; P{TRiP.JF02624}attP2/TM3, Sb1* | BDSC #27315 |
| *UAS-white-RNAi* | *y[1] sc[*] v[1] sev[21]; P{y[+t7.7] v[+t1.8]=TRiP.GL00094}attP2* | BDSC #35573 |
| *Halo::Mam* | *TI{TI}mamRA[Halo]* | DeHaro-Arbona et al., 2024 |
| *GFP::Mam* | *TI{TI}mamRA[sfGFP]* | DeHaro-Arbona et al., 2024 |
| *hnt-MS2* | *$y^{1}$M{vas-Cas9}ZH-2A$w^{118}$, 24xMS2-hnt;* | Hoppe et al. 2020 |
| *E(spl)m7-MS2* | *TI{24xMS2-lacZ-SV40}E(spl)m7-HLH* | Santa-Cruz Mateos et al., 2026 |
| *E(spl)mβ-MS2* | *TI{24xMS2-lacZ-SV40}E(spl)mβ-HLH* | Boukhatmi et al., 2020 |
| *hsp83-MCP::GFP* | *P{w[+mC]=Hsp83-MCP-GFP}3* | BDSC #7280 |
| *His2Av-RFP* | *P{His2Av-mRFP}* | BDSC #23650 |
| *tj-GAL4* | *w[*], P{w[+]=GawB}Tj* | St Johnston lab |
| *hnt$^{peb1}$:GFP* | *w[*], P{w[+]hnt$^{peb1}$-hsp70>GFP}* | Terriente-Félix et al. 2013 |
| *hnt$^{peb2}$:GFP* | *w[*], P{w[+]hnt$^{peb2}$-hsp70>GFP}* | Terriente-Félix et al. 2013 |
| *hnt$^{peb3}$:GFP* | *w[*], P{w[+]hnt$^{peb3}$-hsp70>GFP* | Terriente-Félix et al. 2013 |
| *UAS-Broad-RNAi* | *w[*]; P{w[+mC]=UAS-br.RNAi.R}3* | BDSC #51189 |
| *UAS-mirr-RNAi* | *y[1] sc[*] v[1] sev[21]; P{y[+t7.7] v[+t1.8]=TRiP.HMC06139}attP2* | BDSC #65877 |
| *hnt-[intA]* | *TI{*intA-dsRED*}hnt* | This study |
| *UAS-ParA1-mCherry* | *{w[+] UAS-ParA1-mCherry }attP86Fb* | Gomez-Lamarca et al. 2018 |
| *BrZ1::GFP* | *w[1118]; PBac{y[+mDint2] w[+mC]=br.Z1-GFP.FPTB}VK00033* | BDSC #50754 |
| *UAS-BrZ1* | *w[1118]; P{w[+mC]=UAS-br.Z1}18-13* | BDSC #51190 |
| *1151-Gal4* | *Dmel\P{GAL4}1151* | FBti0007229 |

**Supplementary Table S2: Detailed genotypes for each Figure.**

| Figure | X chromosome | II chromosome | III chromosome |
|---|---|---|---|
| Figure 1B, C | *w[1118]* | | |
| Figure 1D | *w* | *tj-GAL4::tub-Gal80$^{ts}$* | *UAS-LacZ* |
| Figure 1D | *w* | *tj-GAL4::tub-Gal80$^{ts}$* | *UAS-NΔECD* |
| Figure 1E,F | *w* | *tj-GAL4::tub-Gal80$^{ts}$* | *UAS-white-RNAi* |
| Figure 1E,F | *w* | *tj-GAL4::tub-Gal80$^{ts}$* | *UAS-Hairless-RNAi* |
| Figure 2B-H | *hnt-MS2* | *hsp83-MCP::GFP* | *His2Av-RFP* |
| Figure 3B | *w* | - | *hnt$^{peb3}$:GFP* |
| Figure 3C,D,E | *w* | *tj-GAL4::tub-Gal80$^{ts}$* | *UAS-white-RNAi* |
| Figure 3C,D,E | *w* | *tj-GAL4::tub-Gal80$^{ts}$* | *UAS-Broad-RNAi* |
| Figure 3F,G | *w* | *tj-GAL4::tub-Gal80$^{ts}$* | *UAS-white-RNAi/ hnt$^{peb3}$:GFP* |
| Figure 3F,G | *w* | *tj-GAL4::tub-Gal80$^{ts}$* | *UAS-Broad-RNAi / hnt$^{peb3}$:GFP* |
| Figure 4B,C,D | *w/ hnt-[intA]* | *Halo::Mam/ tj-GAL4* | *UAS-ParA1-mCherry* |
| Figure 4B,C,D | *w/ hnt-[intA]* | *tj-Gal4* | *UAS-ParA1-mCherry/ BrZ1::GFP* |
| Figure 5A,B | *hnt-MS2/ hnt-[intA]* | *Halo::Mam/ tj-GAL4:hsp83-MCP::GFP* | *UAS-ParA1-mCherry* |
| Figure 5C,D | *w/ hnt-[intA]* | *tj-Gal4* | *UAS-ParA1-mCherry/ BrZ1::GFP* |
| Figure 5F,G | *1151-Gal4* | *GFP::Mam* | *UAS-NΔECD* |
| Figure 5F,G | w | | *BrZ1::GFP* |

| | | | |
|---|---|---|---|
| Figure 6A,B | *w* | *tj-GAL4::tub-Gal80$^{ts}$* | *UAS-LacZ* |
| Figure 6A,B | *w* | *tj-GAL4::tub-Gal80$^{ts}$/ UAS-BrZ1* | - |
| Figure 6C,D | *w/ hnt-[intA]* | *Halo::Mam/ tj-Gal4* | *UAS-ParA1-mCherry/ BrZ1::GFP* |
| Figure S1A | *w[1118]* | | |
| Figure S1B | *w* | *tj-GAL4::tub-Gal80$^{ts}$* | *UAS-LacZ* |
| Figure S1B | *w* | *tj-GAL4::tub-Gal80$^{ts}$* | *UAS-NΔECD* |
| Figure S1C-F | *w* | *tj-GAL4::tub-Gal80$^{ts}$* | *UAS-white-RNAi* |
| Figure S1C-F | *w* | *tj-GAL4::tub-Gal80$^{ts}$* | *UAS-Hairless-RNAi* |
| Figure S2A | w | *hsp83-MCP::GFP* | *E(spl)m7-MS2/ His2Av-RFP* |
| Figure S2A | w | *hsp83-MCP::GFP* | *E(spl)mβ-MS2/ His2Av-RFP* |
| Figure S2A | *hnt-MS2* | *hsp83-MCP::GFP* | *His2Av-RFP* |
| Figure S2B,C | *hnt-MS2* | *hsp83-MCP::GFP* | *His2Av-RFP* |
| Figure S3A | *w* | - | *hnt$^{peb-1}$:GFP* |
| Figure S3A | *w* | - | *hnt$^{peb-2}$:GFP* |
| Figure S3B | *w* | *tj-GAL4::tub-Gal80$^{ts}$* | *UAS-white-RNAi/ BrZ1::GFP* |
| Figure S3B | *w* | *tj-GAL4::tub-Gal80$^{ts}$* | *UAS-Broad-RNAi/ BrZ1::GFP* |
| Figure S3C | *w* | *tj-GAL4::tub-Gal80$^{ts}$* | *UAS-white-RNAi* |
| Figure S3C | *w* | *tj-GAL4::tub-Gal80$^{ts}$* | *UAS-Broad-RNAi* |
| Figure S3D | *w* | *tj-GAL4::tub-Gal80$^{ts}$* | *UAS-white-RNAi* |
| Figure S3D | *w* | *tj-GAL4::tub-Gal80$^{ts}$* | *UAS-mirr-RNAi* |

| | | | |
|---|---|---|---|
| Figure S4A | *w/ hnt-[intA]* | *Halo::Mam/ tj-GAL4* | *UAS-ParA1-mCherry* |
| Figure S4B | *hnt-MS2/ hnt-[intA]* | *Halo::Mam/ tj-GAL4:hsp83-MCP::GFP* | *UAS-ParA1-mCherry* |
| Figure S4C | *w* | *tj-GAL4::tub-Gal80$^{ts}$ / UAS-Broad(II)* | *hnt$^{peb3}$:GFP* |

**Supplementary Table S3: results of statistical analysis of data presented in Figures**

| Figure | Experiment | n | P Value |
|---|---|---|---|
| Figure 1B, C | *hnt* and *E(spl)*-C genes throughout development | *n=4-25* | - |
| Figure 1D | Notch overexpression in *E(spl)-m3* at stage 9. | n= 10, 6 | *p* = 0.0002498 |
| Figure 1E | Hairless-RNAi in *E(spl)m3* | *n*=5-10 | *p*= 0.0456551<br>*p*= 0.04925248<br>*p*=0.7551 |
| Figure 1F | Hairless-RNAi in *hnt* | n=11-14 | *p*= 0.00018762<br>*p*= 0.0152254<br>*p*=0.20516269 |
| Figure 2D | Active Nuclei Proportion St6 vs St9 | *n=1150,1753 from 5 e.c each* | *p*=1.1353E-118 |
| Figure 2F | Active Periods St6 vs St9 | *n=994,1648 from 5 e.c each* | *p*=1.30163E-25 |
| Figure 2G | Average Maximum Amplitude St6 vs St9 | *n=993 and 1644 nuclei from 5 e.c.* | *p*=0.02577481 |
| Figure 3D | *hnt* in Broad-RNAi | *n=4-7* | *p*= 0.06070842<br>*p*= 0.02484621<br>*p*=0.0080952 |
| Figure 3E | *hnt* ATSs intensities in Broad-RNAi | *n=102, 35, 294, 179, 418, 279* | *p*= 0.12997083<br>*p*= 8.9314E-05<br>*p*=0.00181812 |
| Figure 3F,G | $hnt^{peb-3}$*:GFP* intensities in Broad-RNAi | *n=14,12* | *p*=0.4559 |
| Figure S3B | Validation efficiency of Broad RNAi knockdown | n=120,120, 90, 90 from at least 3 e.c. | *p*= 6.1894E-24<br>*p*=1.5452E-09 |
| Figure S3C | *E(spl)-m7* in Broad RNAi | n=4-7 | *p*= 0.43024168<br>*p*= 0.87066027<br>*p*=0.4215646 |
| Figure S3D | *hnt* in Mirror RNAi | n=11-18 | *p*= 0.467697767<br>*p*= 0.110848273<br>*p*=0.065511726 |
| Figure 4D | Mam at *hnt* locus throughout development | *n=110, 130, 85, 113, 111,100 from at least 3 e.c.* | |
| Figure 4D | Broad at *hnt* locus throughout development | *n=98, 92, 183, 214 from at least 3 e.c.* | |

| | | | |
|---|---|---|---|
| Figure S4A | Percentages of Mam hubs during development | *n=110, 130, 85, 113, 111,100 from at least 3 e.c* | |
| Figure 5A,B and S5A | Mam at *hnt* locus and *hnt*-MS2 movies | *n =12 from 5 e.c* | |
| Figure 5C | Broad at *hnt* locus movies | *n =12 from 5 e.c* | |
| Figure 5D | Hub minimal continuous detection | *n=18,13* | *p*=0.0018247 |
| Figure 5G | FRAP of Mam and Broad | *n=26, 28 from at least 5 animals* | |
| Figure S4B | Cross-correlations Mam vs *hnt*-MS2 | *n =12 from 5 e.c* | |
| Figure 6B | *hnt* transcription in Broad overexpression | *n=7, 20,10* | *p*=0.3474<br>*p*=0.0099 |
| Figure 6C,D | Mam and Broad at *hnt* locus | *n=229 from 9 e.c.* | |
